# Action planning modulates perceptual confidence through action monitoring processes

**DOI:** 10.1101/2023.08.14.553210

**Authors:** Rémi Sanchez, Karen Davranche, Thibaut Gajdos, Andrea Desantis

## Abstract

Dominant models of metacognition suggest that sensory information quality determines perceptual confidence, but recent accounts propose that motor signals also affect confidence judgments. In this study, we investigated the impact of motor planning of perceptual responses on decision confidence, testing two hypotheses. The “fluency hypothesis” suggests that ease of motor response selection and preparation enhances confidence. In contrast, the “monitoring hypothesis” posits that increased action monitoring during response selection boosts confidence, potentially counteracting response fluency. In three pre-registered experiments, participants reported the orientation of a stimulus and indicated their confidence in their response. A cue induced action planning that was either congruent or incongruent with the response side used to report the stimulus orientation. Across experiments, we consistently observed higher confidence when participants prepared spatially incongruent actions compared to congruent ones, regardless of response accuracy. In the third experiment, electroencephalography (EEG) revealed an increased fronto-central P2 amplitude for incongruent actions, suggesting that incongruent action planning heightened early attentional resources needed to resolve response conflict. Incongruent action plans also modulated post-response ERPs at centro-parietal channels (e.g., Pz), typically linked to confidence and error monitoring. These findings align with the “monitoring hypothesis” suggesting that the degree of action monitoring during response selection modulates retrospective decision confidence.

**Public Significance Statement:** While virtually every decision we make leads to an action, the role of motor processes in decision making has been largely neglected. Our results show that retrospective confidence in a perceptual discrimination task is boosted when the motor execution is spatially incongruent with motor preparation, independently of the correctness of the response. Electroencephalography recordings indicate that this effect could be explained by a larger involvement of early attentional resources related to action monitoring, which has an impact on confidence computations. Taken together, these results suggest that action planning information might trigger monitoring mechanisms susceptible to alter retrospective confidence in our decisions, implying that motor processes are not only the output, but also an input of the decision mechanisms.

## Introduction

In a world where feedback is sparse and uncertain, adaptive behavior crucially relies on our ability to estimate how confident we are in the accuracy of our perceptual representations and decisions (Mamassian, 2016). Research showed that the sense of confidence plays a prominent role in learning (Guggenmos et al., 2016), information seeking (Desender et al. 2018), change of mind (Rollwage et al. 2020; Sanchez et al. 2024), and social interactions (Bahrami et al. 2010). Delineating the factors influencing the formation of confidence is therefore crucial for the understanding of adaptive behavior (Rahnev et al., 2021).

Perceptual confidence is typically measured and dissociated from accuracy by asking participants to provide two types of responses during a perceptual decision task. First, they make a perceptual decision about a stimulus, and then they rate their confidence in how accurate that decision was, using a scale (cf. Mamassian, 2016). Classical models of metacognition argue that, just like accuracy, perceptual confidence is determined by the quality of sensory information -i.e., confidence increases with stronger perceptual evidence (Kiani & Shadlen, 2009). In other words, confidence and accuracy are two different descriptions of the same internal variable. However, recent studies suggest that the characteristics of the action used to bring about a perceptual decision can also shape perceptual confidence (Fleming et al., 2015; Gajdos et al., 2019; Pereira et al., 2020; Siedlecka, Koculak, & Paulewicz, 2021; Siedlecka, Paulewicz, & Koculak, 2020; Turner et al., 2021; Wokke, Achoui, & Cleeremans, 2020). Hence, besides sensory evidence, motor signals can modulate confidence judgments. However, it remains unclear what specific aspect of motor processing (i.e., motor preparation or motor execution processes) is responsible for this modulation, and how this modulation takes place.

Two alternative hypotheses, each providing contradictory predictions, can be proposed regarding how motor information is used for confidence estimations. The “fluency hypothesis” suggests that the ease and smoothness in the selection and execution of a motor response, may serve as a cue for confidence judgments (Fleming et al., 2015; see also Wenke et al., 2010; Oppenheimer, 2008; Brouillet et al., 2022; 2023; for notions related to fluency and action fluency). Specifically, fluent actions may lead to higher confidence judgments compared to non-fluent actions. This hypothesis is supported by a study conducted by Fleming et al. (2015), where transcranial magnetic stimulation (TMS) was used to manipulate motor representations in the premotor cortex. The authors found that stimulating premotor areas linked to the chosen perceptual response increased confidence. They hypothesized that this stimulation enhanced action fluency, which, in turn, boosted confidence judgments. The notion of fluency aligns also with the observation that, other things being equal, faster responses are associated with higher confidence reports (Kiani et al., 2014). In sum, the idea underlying this hypothesis is that confidence judgments rely in part on heuristics employing proxies of response accuracy (Ackerman, 2019; Maniscalco et al., 2016; Turner et al., 2021; Van Marcke et al., 2022), and action fluency would be one of these proxies. Alternatively, according to the “monitoring hypothesis”, the modulation of action signals on confidence occurs through the involvement of action monitoring processes (cf. Anzulewicz et al., 2019; Gajdos et al., 2019; Sanchez et al., 2024). Contrary to the fluency hypothesis, this perspective suggests that it is not the fluency of an action that leads to higher confidence estimates, but rather the degree of control exerted and the level of monitoring required for implementing a perceptual decision. The higher the control required for implementing a response, the higher the confidence in the perceptual decision. This hypothesis is supported by neuroimaging research highlighting the critical role of the Prefrontal Cortex (PFC) in both perceptual confidence and in the monitoring of self-generated decisions (Fleming et al., 2012), as well as by more recent studies. For instance, Turner et al. (2021) showed that effortful perceptual responses are associated with higher confidence than less effortful motor decisions (but see Hagura et al. 2023). Similarly, Sanchez et al. (2024) showed that enhanced visuo-motor control towards perceptual decisions increase confidence judgments. Finally, Gajdos et al. (2019) observed that when undesired motor responses are correctly inhibited perceptual confidence increases. Overall, these studies suggest that when action monitoring processes are more strongly involved - either because actions are more effortful or unwanted responses need to be controlled and inhibited (Anzulewicz et al., 2019; Gajdos et al., 2019; Morel et al., 2017; Sanchez et al., 2024; Turner et al., 2021)-confidence increases, even if action fluency is hindered.

To confront these two hypotheses and test their predictions, we conducted three pre-registered experiments which aimed specifically at investigating how motor preparatory signals of perceptual responses are used in confidence estimations. Specifically, participants reported the orientation of a Gabor (vertical or horizontal) with their index fingers and judged their confidence in their decision. Using a novel motor priming paradigm, we prompted participants to prepare an action in advance that could either be compatible or incompatible with the action ultimately chosen to report the Gabor orientation.

The “fluency hypothesis” would predict that perceptual confidence increases when the perceptual response is compatible with the primed action. Indeed, priming a compatible aspect of the perceptual response should facilitate the planning and selection of that response, thereby increasing fluency (e.g., Chambon & Haggard 2012). Thus, higher confidence is expected when participants prepare a congruent action (e.g., a left response is primed and the Gabor is reported with a left response) compared to incongruent actions (e.g., a left response is primed but the Gabor is reported with a right response). In contrast, the “monitoring hypothesis” would predict the opposite: perceptual confidence is higher when the primed action is incompatible with the subsequent perceptual response. In incompatible trials, increased monitoring is required to control and inhibit unwanted motor plans. Accordingly, higher confidence may be expected when participants prepare an incongruent action compared to congruent actions.

In Experiment 3, we used electroencephalography (EEG) to investigate the specific brain processes modulating confidence, focusing in particular on the frontocentral P2/N2 complex. The P2 component is linked to attentional resource allocation for feature-based stimulus evaluation and response selection (Darriba et al., 2018; Xie et al., 2020), while the N2 is associated with motor inhibition and it is typically observed in go/no-go tasks (Folstein et al., 2008). We predicted that priming incompatible responses would require higher monitoring and control, leading to increased P2 and N2 components.

In a nutshell, we found that confidence was higher when participants prepared incongruent rather than congruent actions, supporting the monitoring hypothesis over the fluency hypothesis: the more individuals monitor and control their motor performance, the higher their perceptual confidence. This finding is important as it suggests that action processes are not merely auxiliary to decision-making; instead, they shape high-level cognitive constructs, such as retrospective confidence.

## Materials and Methods

### Materials

*Participants.* For Experiment 1, data collection was stopped after gathering the data from 16 participants that did not violate our data inclusion criteria. The inclusion criteria required participants to achieve perceptual performance levels suitable for the study of confidence (i.e., an accuracy rate between 60% and 90%), and to use each level of confidence at least once, indicating a proper understanding of the task (for a complete description of the inclusion criteria, see Supplementary Information -SI-, section *Experimental Procedure*). Crucially, we verified that our results remained consistent with both the inclusion and exclusion of outlier participants (SI *Table S6, S7 & S8*). This sample size was estimated by performing a statistical power analysis (using G*Power; Faul et al., 2007), based on a pilot study that investigated the effect of our motor priming paradigm on response times (see OSF pre-registration: MG1). Ten participants were not included in the sample since they did not fulfil our inclusion criteria. The remaining 16 participants (11 participants reported their gender as female, 5 as male, mean age = 24.9, SD = 4.18) were analyzed. Based on the results of the first experiment (see OSF pre-registration: MG2) and on the decision to increase statistical power, the data collection of Experiment 2 was stopped after gathering the data from 24 participants. In total 32 adults were recruited for this experiment, 8 of them were not included in the sample since they did not fulfil our inclusion criteria. The remaining 24 participants (14 participants reported their gender as female, 10 as male, average age = 25.8, SD = 3.68) were analyzed. For Experiment 3, we replicated the procedure of Experiment 2 (see OSF pre-registration: MG3) and no participant was excluded, hence 24 participants (12 participants reported their gender as female, 12 as male, average age = 25.4, SD = 4.36) were recruited for this experiment and included in the analyses. Participants were paid 10€/hour in gift card (and 15€/hour for Experiment 3 since it involved EEG). All participants had normal or corrected-to-normal vision, were right-handed, and were naïve regarding the hypothesis under investigation. They all gave written and informed consent before participating in the experiment. This study was conducted in agreement with the requirements of the Declaration of Helsinki and approved by the ethics committee of Université Paris Cité (Number: 00012024-51).

*Equipment.* Participants performed the experiment in a dark room. They were comfortably seated in front of an LCD monitor (in Experiments 1 and 2 we used a 20-inch Asus PG248Q with 1920 by 1080 screen resolution and a refresh rate of 60 Hz, in Experiment 3 we used Display ++ with a resolution of 1920 x 1080 pixels, a size of 32 inches, and refresh rate of 60 Hz). The experiment was programmed with Python (Python Software Foundation. Python Language Reference, version 3.9. Available at http://www.python.org) and the stimuli were generated using the Psychopy software (Peirce et al., 2019). A button response box (Millikey Response Box MH-5) and foot pedals (Accuratus X3P Footswitch) were used to collect participants’ hand and feet responses, respectively. For Experiment 3, two thumb-press buttons were specifically built for convenience (See SI, *Figure S1*). EEG signals were recorded with 64 Ag/AgC1 electrodes mounted on an elastic cap and amplified by an ActiCHamp Plus amplifier (Brain Products GmbH, Munich, Germany). Electrodes were arranged according to the international 10–20 systems. EEG data were online referenced to M2. EEG signal was sampled at a digitization rate of 500 Hz.

### Methods

*Experimental paradigm.* In the three experiments, we manipulated the motor preparation of perceptual responses, by randomly interleaving the trials of a perceptual task with the trials of a speeded Reaction Time (RT) task. In each trial (of both the perceptual and the speeded RT task) a visual cue was displayed (e.g., an arrow in Experiment 2 and 3, see below). The visual cue indicated the action to perform (e.g., left or right button press) in the speeded RT task. This action had to be executed as soon as a white flash was presented. However, in part of the trials, participants rather than viewing a white flash, were presented with a Gabor at perceptual discrimination threshold. Thresholds were calculated for each participant in a preliminary calibration phase (see Calibration Phase in the Supplementary Material). In those trials, they were instructed to ignore the speeded RT task and to perform instead a perceptual task, which consisted in indicating the orientation of the Gabor.

Crucially, the task to perform at each trial was not known at the time of the cue onset, only the presentation of the white-flash or the Gabor indicated the participant which task to perform. We expected that the presentation of the visual cue would push the participants to prepare in advance a response in order to meet the temporal constraint imposed by the speeded RT task. The action preparation induced by the cue would then facilitate (or interfere with) the selection/execution of a compatible (or incompatible) action required to report the orientation of the Gabor.

### Experiment 1

Each trial started with a presentation of a green fixation cross (0.4 by 0.4 degrees of visual angle, dva) at the center of the screen for a duration of 300 ms (see Figure 1). A visual cue was then displayed for 200 ms. The cue was the same fixation cross but presented with a different color (e.g., blue or yellow). In the speeded RT task trials participants were instructed to report the color of the cue by executing a left or a right response as soon as a white flash (1.8 dva diameter) was displayed (e.g., left-response if the fixation turned blue and right-response if the fixation turned yellow). In half of the trials, participants reported the color of the fixation with their index fingers (the same pair of effectors used in the perceptual task described below, i.e., same effector block), while in the other half of the trials, they reported the color by using their feet (different effector block). The white flash was presented 800 ms after the onset of the cue and displayed for a duration of 50 ms. Participants had 800 ms to respond to the cue. If no response was given within this delay, the trial was interrupted and a message ‘too late!’ was displayed. The action-color mapping and the order of the blocks (same and different effector) were counterbalanced across participants.

**Figure 1.**
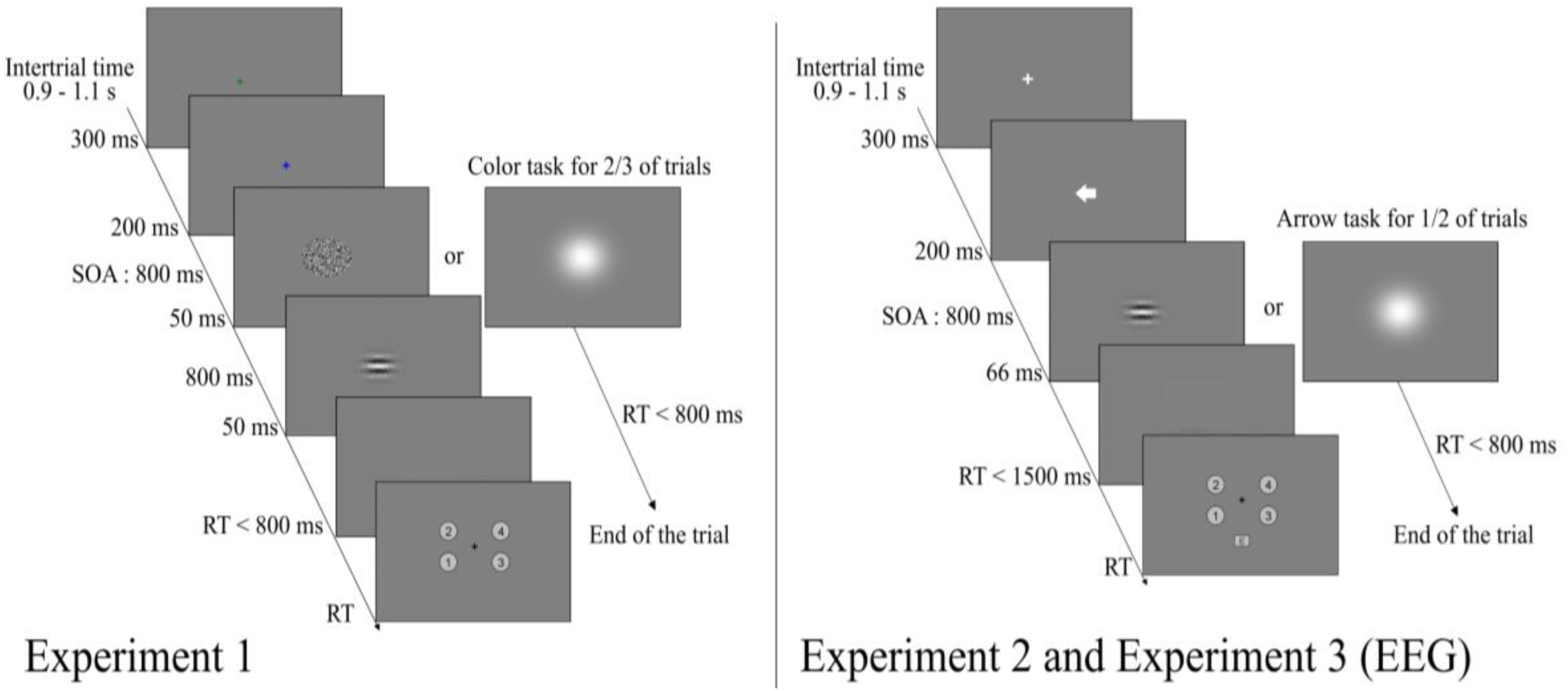
Experimental paradigms. Participants are initially presented with a visual cue (i.e., a colored fixation in Experiment 1, and an arrow in Experiments 2 and 3). Subsequently, they view either a white flash or a Gabor patch. If a white flash is displayed, participants have to report the color of the visual cue (Experiment 1) or its pointing direction (Experiment 2 and 3) by pressing a left or a right button (speeded RT task). If a Gabor is presented, they have to report its orientation by executing a left or a right button-press (e.g., left for horizontal and right for vertical; perceptual task). After their perceptual decision, participants report their confidence on a scale from 1 to 4. For Experiment 2 and 3, participants could also report an error instead of a confidence level.

In one third of the trials, rather than viewing a white flash, participants were presented with a noise texture (1.8 dva diameter) for 50 ms. A Gabor patch (1.8 dva diameter) was then displayed 800 ms after the onset of the noise stimulus, for a duration of 50 ms. This delay of 800 ms was introduced in Experiment 1 to help participants to prepare for reporting the orientation of the Gabor. In these trials, participants were instructed to forget about the speeded RT task and were asked to perform a **perceptual task** instead. Importantly, perceptual and speeded RT trials were randomly interleaved. The perceptual task consisted in discriminating the orientation (horizontal or vertical) of the Gabor patch (see Figure 1). The Gabor was embedded in a noisy texture and displayed with a contrast level supporting 72.5% correct discrimination performance calculated separately for each participant in a preliminary calibration phase (see SI *Experimental procedure*). Perceptual responses were provided by executing a left or right index finger key-press as quickly and as accurately as possible (Gabor-response mapping was counterbalanced across participants). Participants had 800 ms to respond. If no response was given within this delay, the trial was interrupted with an error message and added to the trial list.

Critically, the action performed to report the orientation of the Gabor could be compatible or incompatible with the action prepared to report the visual cue. In particular, we manipulated two sorts of compatibility. Firstly, the action prepared to report the cue could be spatially congruent or incongruent with the response selected to report the orientation of the Gabor. For instance, in spatially incongruent trials, the cue induced the preparation of a left (index finger or foot) response while the participant performed a right (index finger) response to report the Gabor. Secondly, the action prepared to report the cue could involve the same (index fingers) or a different pair of effectors (feet) than those used by the participant to report the Gabor. Hence, the experiment implemented a 2 by 2 factorial design with *Spatial congruency* (congruent and incongruent) and *Effector compatibility* (same and different).

After the perceptual response, participants reported their confidence on a scale from 1 to 4 corresponding to the probability that their perceptual decision was correct (1: guessing, 2: barely confident, 3: fairly confident, 4: certainly correct). The confidence level numbers were displayed inside four circles randomly displayed around the center of the screen (Figure 1). There was no time pressure to report confidence ratings. To indicate the level of confidence at each trial, since participants used four effectors in the experiment, each of the four effectors were associated with a confidence circle.

The experiment lasted about one hour and thirty minutes and was divided into two main blocks (same and different effector blocks). Each block was composed of 11 mini-blocks of 36 trials each (24 speeded RT task trials and 12 perceptual task trials) for a total of 792 trials evenly split into 22 blocks. Before the main experiment, a training phase and a calibration phase were performed (see SI *Experimental procedure*).

### Experiment 2

For Experiment 2, we made a few changes to the design of Experiment 1. In order to make the speeded RT task easier and more intuitive, the visual cue was a white arrow (1 dva by 1 dva) displayed at the center of the screen pointing either to the left or to the right. Participants were instructed to indicate its direction with a left or right response respectively (Figure 1). Furthermore, in the *different effector* block we replaced left and right foot responses with left and right middle finger button-presses. We changed the effectors because in Experiment 1 we observed a clear deterioration of performances when switching from lower- to upper-limb actions. Since the speeded RT task was easier, it enabled us to bring the onset of the Gabor patch closer to the cue. Hence, we removed the warning noise texture presented before the occurrence of the Gabor, which was now displayed 800 ms after the cue (thus, respecting the timing of the presentation of the visual-flash in the speeded RT task).

We also increased the response time limit of the perceptual task to 1500 ms. Additionally, in case participants made an involuntary mistake in the perceptual task (e.g., they executed a right key-press but they wanted to perform a left key-press), they could report it during the confidence judgment by pressing the space bar (Figure 1).

The experiment lasted ∼45 minutes and was divided into two main blocks (same effector and different effector). Each block was composed of 5 mini-blocks of 40 trials each (20 cue and 20 Gabor trials) for a total of 400 trials evenly split into 10 blocks.

### Experiment 3

In Experiment 3, the design was identical to Experiment 2, except for the following changes. Since effector compatibility did not seem to impact confidence, we removed the different effector condition, so the speeded RT task and the perceptual task were performed with the thumbs. Participants indicated their confidence by moving an arrow pointing towards one of the confidence levels. Participants could move the arrow either clockwise (right hand) or anti-clockwise (left hand) and validate their confidence level with a double button-press. To report an error, participants had to validate when the arrow was pointing toward the box ‘E’ (Figure 1). Additionally, the confidence display was presented 1.25 seconds after the perceptual response to leave some time without visual stimulation for EEG considerations. The experiment was divided into 10 mini-blocks of 40 trials each (20 cue and 20 Gabor trials) for a total of 400 trials lasting about 45 minutes.

### Data analyses

**Behavioral data.** Data were analyzed using R (R Core Team, 2020). We used generalized linear mixed-effect models (inverse-gaussian with identity log function) for Response Times (RT), generalized linear mixed-effects models (binomial with logit link function) for response accuracy, and ordinal regression with cumulative link model for confidence. Linear and generalized linear regressions were performed with the package lme4 (Bates et al., 2015) and ordinal regressions with the package ordinal (Christensen et al., 2018). Linear regressions were performed with the restricted maximum likelihood fitting method, and *p*-values for coefficients were computed with Satterthwaite’s method using the lmerTest package (Kuznetsova et al., 2017). Ordinal regressions were performed with Laplace approximation, and *p*-values were computed for Wald tests. The models were adjusted to accommodate convergence or singularity issues. The structure of random effects was determined based on the parsimony principle (Bates et al., 2015). Notably, for some analyses, the inclusion of the maximal random structure led to convergence failures. In those cases, we performed a Principal Component Analysis (PCA) to isolate the random effects that the least contributed to model fitting and we removed them one-by-one from the final model, until there were no convergence failures. We used an alpha level of 0.05 for all statistical tests. Factors were coded using sum contrasts, and response times were centered (with respect to the across-subjects mean). In addition, our results were replicated with the inclusion of outliers (see SI *Tables S6, S7 & S8*), and they were also replicated using Bayesian methods (see SI *Bayesian analyses, Tables S3, S4 & S5*). For graphical representations, the package ggplot2 was used (Wickham, 2009).

In Experiment 2 and 3, participants were able to report errors they made consciously when indicating the orientation of the Gabor. These trials were omitted from further analysis. In Experiment 2 they represented 6.0 % of the total number of trials (i.e., 290 errors over 4800 trials in total). Specifically, participants reported 151 errors in the different effector condition (44 in the incongruent and 107 in the congruent condition) and 139 in the same effector condition (52 in the incongruent and 87 in the congruent condition). In Experiment 3 they represented 5.0 % of the trials (i.e., 230 error over 4800 trials in total). Specifically, participants reported 133 errors for congruent and 74 for incongruent trials.

**EEG Data.** EEG data were analyzed using MNE-Python 1.3.0 (Gramfort et al., 2013, 2014) and re-referenced to the left and right mastoids (M1 and M2). The signal was band-pass filtered between 0.05 and 48 Hz by a non-causal infinite impulse response filter. The raw data was inspected visually to remove time periods containing large artifacts. Subsequently, we performed an independent component analysis (Hyvärinen, 1999), to identify and remove components representing blinks or eye movements. We then extracted epochs going from −1900 ms to 2500 ms time-locked to Gabor onset. A second artifact rejection was performed. Epochs containing amplitudes greater than 100 µV or less than −100 µV were marked as potential artifacts and then removed after confirmation through visual inspection. This led to the removal of 224 out of 4628 trials, i.e., a proportion of 0.048 trials. EEG activity in specific time windows for ERP components was quantified through mean amplitudes calculation and tested with ANOVA and t-tests as they are robust to different numbers of trials across conditions (Luck, 2012). If no specific time windows were identified from the literature for a component, we performed a nonparametric cluster-based permutation test (Maris and Oostenveld, 2007) to isolate the time windows exhibiting a difference between congruent and incongruent trials.

Firstly, we performed a temporal cluster-based permutation test comparing the voltage amplitude observed in congruent and incongruent trials on a time window going from 0 to 350 ms post-stimulus (we selected a time window in which no perceptual response could have already occurred) at electrode Fz, Cz and FCz. Stimulus-locked segments were corrected with a baseline of 200 ms prior to Gabor onset (Folstein & Van Petten, 2008; Luck et al., 2012). This analysis was conducted to identify the time window and potential ERP component (e.g., P2 or N2) that differed between congruent and incongruent trials.

Secondly, we analyzed post-response ERPs. Response-locked epochs were corrected with a 100 ms baseline, going from −100 to 0 ms prior to the response (Boldt & Yeung., 2015). We focused and analyzed a time period going from 0-500 ms post-response at the electrode Pz, given that this time window and electrode were already identified in previous study of EEG correlates of confidence (Davies et al., 2001; Wang et al., 2020; Rausch et al., 2020). We then performed a cluster-based permutation tests on the difference between congruent and incongruent between 0 ms to 500 ms post-response at Pz to identify potential components exhibiting a modulation of spatial congruency. Two temporal clusters were identified (see Results). To investigate their relation with confidence judgments we binned the four levels of confidence into low (confidence ratings equal 1 or 2) and high (confidence ratings equal 3 or 4) trials and performed ANOVAs on the by-participant average amplitude observed in the two time periods reported above. The ANOVA included Confidence (high, low) and Congruency (congruent, incongruent) as factors. Note that we also analyzed the amplitude of post-stimulus frontal theta oscillations. These analyses did not yield conclusive results and are reported and discussed in the supplementary materials.

Furthermore, we analyzed the Lateralized Readiness Potential (LRP), traditionally associated with motor preparation (Coles, 1989). Our motor priming paradigm aimed to induce the preparation of a specific action prior to the perceptual response. Hence, we expected shorter response times for congruent compared to incongruent actions, along with similar changes in the latency of congruent LRPs. LRPs were calculated using the double subtraction method, where the average effector-specific activity over the left and right motor cortices (electrodes C3 and C4) are subtracted (Coles, 1989). Response-locked LRP epochs (going from −800 to 0 ms time-locked to perceptual response onset, Smulders & Miller, 2012) were corrected with a baseline going from −1000 and −800 ms prior to the perceptual response (Kraemer & Gluth, 2023). A jackknife-based method with a one-third threshold was employed to investigate potential differences in LRP onset latencies between congruent and incongruent responses, as this method has been shown to be a powerful tool for analyzing LRP onset latencies (Ulrich & Miller, 2001).

Additional analyses on motor preparatory signals and the P300 were conducted as sanity checks and are reported in the Supplementary Information. Specifically, we examined the relationship between confidence and the stimulus-locked P300 component, given that previous research linked the P300 to evidence accumulation, sensory uncertainty, and confidence judgments (Herding et al., 2019). Our findings indeed revealed a correlation between confidence and the P300 (see SI, EEG analyses and results, Figure S6).

#### Transparency and openness

The hypotheses, methods, and statistical procedure for the behavioral data were pre-registered before the experiment. Pre-registration, data and codes for analysis are publicly available at https://osf.io/sb45z/. However, the analysis of response time was not included in the pre-registration. In addition, we made some adjustments to the pre-registered statistical procedure due to convergence issues with mixed models (see Methods). In particular, for the analysis of confidence, in Experiment 1, we removed the interaction between congruency and effector from the random effect structure; and in Experiment 3, we removed the interaction between congruency and response time from the random effect structure. For the analysis of accuracy, in Experiment 1, we only kept response time in the random effect structure; in Experiment 2, we only kept response time and effector in the random effect structure; and in Experiment 3, we only kept response time in the random effect structure. Regarding the EEG analyses, only some general aspects were specified in the pre-registration, as the exact signals and methods of investigation were not fully determined at the time. We indicated in the pre-registration that we would examine brain oscillations, such as post-stimulus theta activity (4-7 Hz) in fronto-parietal electrodes, as a marker of cognitive control (Eisma et al. 2021). Although these analyses were performed, they are not included in the manuscript because they did not yield clear results. This is mentioned and discussed in the Supplementary Material.

## Results

### Behavioral Result

The analyses encompassed a total of 4,224 trials for Experiment 1; 4,510 trials for Experiment 2; and 4,570 trials for Experiment 3. The average task accuracy for Experiment 1 (M = 0.68, SE = 0.03), Experiment 2 (M = 0.73, SE = 0.01), and Experiment 3 (M = 0.73, SE = 0.02), and the average confidence ratings for Experiment 1 (M = 2.34, SE = 0.12), Experiment 2 (M = 2.51, SE = 0.13), and Experiment 3 (M = 2.1, SE = 0.7) were appropriate for investigating confidence judgments, aligning with previous studies (Fleming & Lau, 2014; Ferrigno et al. 2019).

### Accuracy

We analyzed accuracy using a binomial generalized linear mixed-effects model, with a logit link function. Statistical significance was assessed with the Wald test. Formally, we estimated the following models:

Experiment 1: accuracy ∼ congruency * effector * effector_order * RT + (1 + RT | subject)

Experiment 2: accuracy ∼ congruency * effector * effector_order * RT + (1 + effector + RT | subject)

Experiment 3: accuracy ∼ congruency * RT + (1 + RT | subject)

Detailed results are gathered in Table S1 in the Supplementary Material. No effect of spatial congruency on accuracy was observed in Experiment 1 (p = 0.91) and Experiment 2 (p = 0.1). Incongruent responses were slightly more accurate (M = 0.74, se = 0.02) than congruent responses (M = 0.72, se = 0.02) in Experiment 3 (odds-ratio = 0.92, p = 0.02). Accuracy was higher in the same effector block compared to the different effector block in Experiment 1 (odds-ratio = 1.1, p < 0.001), with lower accuracy when the visual cue primed feet responses (M = 0.64, se = 0.02) compared to hand responses (M = 0.71, se = 0.01), indicating that the visual task was harder when it required feet responses. This effect was not observed in Experiment 2 (p = 0.69). No interaction between congruency and effector (Experiment 1: p = 0.81; Experiment 2: p = 0.92). In Experiment 1, we observed a significant interaction between effector order and response times on accuracy (odds-ratio = 0.36, p = 0.003, see SI, *Figure S4*), indicating that only for incorrect responses, response times were faster when participants completed the same effector block first compared to when they began with different effector block.

### Response time

Response times (RT) were calculated with respect to the onset of the Gabor patch. They were analyzed with a generalized (inverse-gaussian with identity link function) linear mixed-effects model, and p-values were computed with Satterthwaite’s method.

Experiment1: RT ∼ accuracy * congruency * effector * effector_order + (1 + accuracy + effector | subject)

Experiment 2: RT ∼ accuracy * congruency * effector * effector_order + (1 + accuracy + effector + effector_order | subject)

Experiment 3: RT ∼ accuracy * congruency + (1 + accuracy_gabor + congruency | subject).

Detailed results are gathered in Table S2 in the Supplementary Material. No effect of spatial congruency on RT was observed in Experiment 1 (p = 0.17). This suggests that the presentation of the noise patch (see Method section) 800 ms before the Gabor allowed participants to more easily control primed responses without any impact on response times. However, RTs were faster for congruent (Experiment 2: M = 695 ms, se = 20 ms; Experiment 3: M = 673 ms, se = 23 ms) than incongruent trials (Experiment 2: M = 713 ms, se = 17 ms; Experiment 3: M = 694 ms, se = 23 ms) in Experiment 2 (β = −11.5, p < 0.001) and Experiment 3 (β = −13.7, p = 0.008). This suggests that our action priming successfully facilitated (impeded) the selection of spatially congruent (incongruent) perceptual responses. In Experiment 1, we found an interaction between effector and effector order (β = −31.38, p < 0.001; see SI, Figure S4 and post-hoc tests), indicating that participants were overall faster in the second block. Notably, if they started with the same effector block, their reaction times were faster in the different effector block (i.e., the second block), and those who started with the different effector block showed faster responses in the same effector block (i.e., the second block for those participants; see SI, Figure S4 & Post-hoc tests).

### Confidence

We analyzed confidence using a cumulative link mixed-effects model, with a probit link function. Statistical significance was assessed with the Wald test. Formally, we estimated the following models:

Experiment 1: confidence ∼ accuracy * congruency * effector * effector_order + RT + (1 + accuracy + congruency + effector + effector_order + RT | subject)

Experiment 2: confidence ∼ accuracy * congruency * effector * effector_order + RT + (1 + accuracy + congruency * effector + effector_order + RT | subject)

Experiment 3: confidence ∼ accuracy * congruency + RT + (1 + accuracy + congruency + RT | subject).

Detailed results are gathered in Table 1 below. As expected, confidence was greater for correct trials than for errors in all three experiments (Experiment 1: odds-ratio = 1.9, p < 0.001; Experiment 2: odds-ratio = 1.4, p < 0.001; Experiment 3: odds-ratio = 1.6, p < 0.001). We also found in Experiments 2 and 3 the classical relation between RT and confidence, with faster responses leading to higher confidence levels (Experiment 2: odds-ratio = 0.07, p < 0.001; Experiment 3: odds-ratio = 0.14, p < 0.001). This relation was however not observed in Experiment 1 (p = 0.17). This might be due to a floor effect, response time being much faster in Experiment 1 than in the two others (see Figure 2).

**Figure 2.**
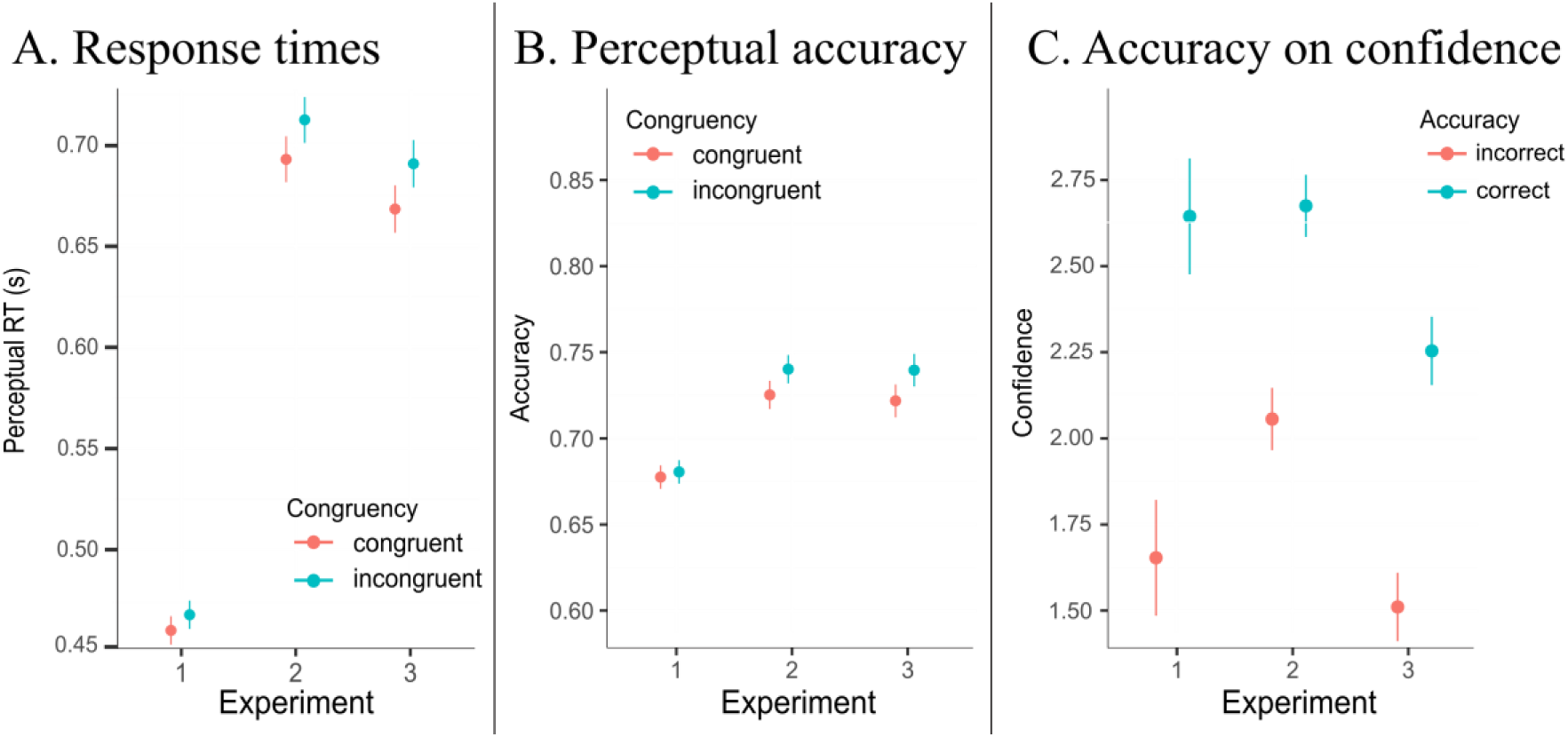
Average response times (A), and average accuracy (B) in the perceptual task, as a function of spatial congruency for each experiment. The graph C depicts perceptual confidence as a function of perceptual accuracy (C). Error bars represent confidence interval regarding the dispersion of individual averages corrected by the Cousineau & Morey method (Morey, 2008).

**Table 1.** Hierarchical regression tables predicting confidence from accuracy, congruency, and reaction time for Experiment 3 (right), plus effector and effector order for Experiment 1 (left), 2 (middle), and 3 (right). Coefficients were coded as follows: Accuracy: error = −1, correct = 1; Congruency: incongruent = −1, congruent = 1; Effector compatibility: different effector (feet in Experiment 1, middle fingers in Experiment 2) = −1, same effector (index fingers) = 1; Bloc order: same effector first = −1, different effector first = 1. Response time is mean-centered. The models tested were the following: Experiment 1: confidence ∼ accuracy * congruency * effector * effector_order + RT + (1 + accuracy + congruency + effector + effector_order + RT | subject) Experiment 2: confidence ∼ accuracy * congruency * effector * effector_order + RT + (1 + accuracy + congruency * effector + effector_order + RT | subject) Experiment 3: confidence ∼ accuracy * congruency + RT + (1 + accuracy + congruency + RT | subject)

| *Predictors* | **Confidence (Exp. 1)** |  |  | **Confidence (Exp. 2)** |  |  | **Confidence (Exp. 3)** |  |  |
| --- | --- | --- | --- | --- | --- | --- | --- | --- | --- |
|  | *Odds Ratios* | *CI* | *p* | *Odds Ratios* | *CI* | *p* | *Odds Ratios* | *CI* | *p* |
| 1\|2 | 0.66 | 0.50 – 0.86 | **0.002** | 0.43 | 0.31 – 0.60 | **<0.001** | 0.85 | 0.74 – 0.99 | **0.037** |
| 2\|3 | 1.66 | 1.27 – 2.17 | **<0.001** | 1.23 | 0.89 – 1.70 | 0.209 | 2.36 | 2.03 – 2.74 | **<0.001** |
| 3\|4 | 4.49 | 3.43 – 5.88 | **<0.001** | 3.12 | 2.25 – 4.32 | **<0.001** | 6.05 | 5.17 – 7.07 | **<0.001** |
| accuracy gabor [1] | 1.90 | 1.64 – 2.20 | **<0.001** | 1.42 | 1.32 – 1.52 | **<0.001** | 1.58 | 1.43 – 1.75 | **<0.001** |
| congruency [1] | 0.94 | 0.90 – 0.98 | **0.004** | 0.94 | 0.89 – 0.99 | **0.017** | 0.90 | 0.86 – 0.94 | **<0.001** |
| effector [1] | 1.01 | 0.94 – 1.09 | 0.781 | 0.98 | 0.88 – 1.09 | 0.668 |  |  |  |
| effector order [1] | 0.78 | 0.61 – 0.99 | **0.043** | 0.74 | 0.54 – 1.03 | 0.074 |  |  |  |
| rt gabor centered | 0.59 | 0.28 – 1.26 | 0.175 | 0.07 | 0.04 – 0.12 | **<0.001** | 0.14 | 0.09 – 0.22 | **<0.001** |
| accuracy gabor [1] * congruency [1] | 1.04 | 1.00 – 1.09 | **0.028** | 1.00 | 0.96 – 1.04 | 0.937 | 1.03 | 0.99 – 1.07 | 0.185 |
| accuracy gabor [1] * effector [1] | 1.08 | 1.04 – 1.13 | **<0.001** | 1.01 | 0.97 – 1.06 | 0.479 |  |  |  |
| congruency [1] * effector [1] | 0.98 | 0.94 – 1.01 | 0.221 | 0.98 | 0.94 – 1.02 | 0.258 |  |  |  |
| accuracy gabor [1] * effector order [1] | 0.94 | 0.81 – 1.09 | 0.396 | 1.03 | 0.96 – 1.10 | 0.444 |  |  |  |
| congruency [1] * effector order [1] | 1.03 | 0.99 – 1.07 | 0.169 | 1.04 | 0.99 – 1.10 | 0.107 |  |  |  |
| effector [1] * effector order [1] | 0.88 | 0.82 – 0.96 | **0.003** | 0.92 | 0.83 – 1.02 | 0.110 |  |  |  |
| (accuracy gabor [1] * congruency [1]) * effector [1] | 1.02 | 0.98 – 1.06 | 0.342 | 1.03 | 0.99 – 1.07 | 0.127 |  |  |  |
| (accuracy gabor [1] * congruency [1]) * effector order [1] | 1.00 | 0.96 – 1.04 | 0.875 | 0.96 | 0.92 – 0.99 | **0.024** |  |  |  |
| (accuracy gabor [1] * effector [1]) * effector order [1] | 1.08 | 1.04 – 1.12 | **<0.001** | 1.00 | 0.96 – 1.04 | 0.926 |  |  |  |
| (congruency [1] * effector [1]) * effector order [1] | 1.00 | 0.96 – 1.04 | 0.924 | 1.03 | 0.98 – 1.07 | 0.252 |  |  |  |
| (accuracy gabor [1] * congruency [1] * effector [1]) * effector order [1] | 1.01 | 0.97 – 1.04 | 0.792 | 0.99 | 0.95 – 1.03 | 0.460 |  |  |  |

There was no main effect of effector compatibility on confidence (Experiment 1: p = 0.78; Experiment 2: p = 0.67), and no significant interaction between spatial congruency and effector compatibility (Experiment 1: p = 0.22; Experiment 2: p = 0.26, see Table 1). We also observed some interactions involving effector compatibility, though these were not consistent across experiments. These interactions are discussed and reported in full in the Supplementary Material. Overall, they suggest that switching from upper to lower limb responses in the speeded RT task of Experiment 1 introduced an attentional cost affecting both confidence and accuracy (i.e., perceptual and metacognitive accuracy decreased in the different effector block). Additionally, effector compatibility interacted with block order. Accordingly, the difficulty of switching from upper to lower limbs varied depending on whether participants began the task with hand or foot responses. These interactions did not seem to be consistent as they were not replicated across experiments.

Importantly, priming the side of an action consistently modulated confidence across the three experiments, over and above the specific effector primed, accuracy and reaction times. Indeed, we observed a main effect of spatial congruency, with higher confidence in incongruent than congruent trials in the three experiments (Experiment 1: odds-ratio = 0.94, p = 0.004; Experiment 2: odds-ratio = 0.94, p = 0.017; Experiment 3: odds-ratio = 0.90, p < 0.001). This effect was slightly larger for errors than for correct trials in Experiment 1 (odds-ratio = 1.04, p = 0.028). However, the interaction between congruency and accuracy was not significant in Experiment 2 (p = 0.94), nor in Experiment 3 (p = 0.18). Only in Experiment 2 we observed a three-way interaction involving accuracy, spatial congruency and effector order (odds-ratio = 0.98, p = 0.024, see SI Figure S3). However, post-hoc tests revealed no significant contrasts (see SI, post-hoc tests), leaving some statistical ambiguity regarding the interpretation of this interaction.

Hence, the consistent finding across experiments is the modulation of spatial congruency on confidence, with higher confidence for spatially incongruent actions compared to spatially congruent ones. Importantly, this effect was observed even after controlling for accuracy and response times in the models, and even when perceptual accuracy did not significantly differ between congruent and incongruent trials (see Experiments 1 and 2). The direction of the congruency effect on confidence supports the action monitoring hypothesis rather than the action fluency hypothesis.

**Figure 3.**
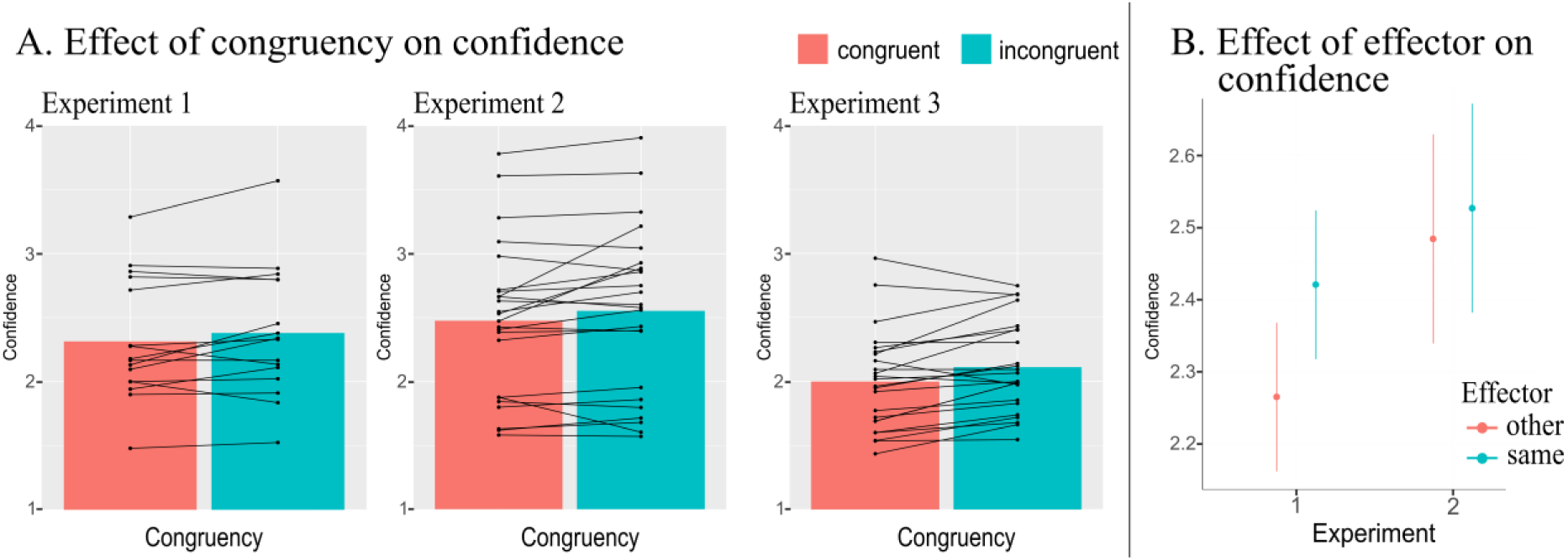
The graphs in A show the across-participant average confidence and individual confidence values as a function of spatial congruency for each experiment. The graph in B shows the across-participant average confidence as a function of effector compatibility for Experiments 1 and 2. Error bars represent confidence intervals indicating the dispersion of individual averages, corrected using the Cousineau & Morey method (Morey, 2008). Note that confidence averages are presented only for illustrative purposes. Statistics were not performed on averages, since confidence were treated as an ordinal variable.

### ERP Results (Experiment 3)

**Post-stimulus activity.** To investigate the way action priming may have influenced confidence judgment we analyzed whether congruency modulated EEG markers of cognitive control that are typically observed after stimulus presentation, such as the P2/N2 complex (and theta oscillations, see SI). We reasoned that the action prepared in response to the visual cue could have conflicted with the action required to report the orientation of the Gabor (i.e., incongruent trials). Specifically, preparing a congruent action in advance may have facilitated response selection, while preparing an incongruent action may have led the participant to control and inhibit unwanted choices as soon as the Gabor was presented. This conflict would be observed rather early after the onset of the Gabor.

A cluster-based permutation test performed on a time window going from 0 ms to 350 ms post-stimulus onset (i.e. we selected a time window in which no perceptual response could have already occurred), identified a significant temporal cluster at FCz (t(23) = −2.6, p < 0.05, Figure 4B) going from 190 to 240 ms (similar results for Fz and Cz, see SI *Figure S7 & S8*), i.e., in a time period where the P2 component is typically observed (Ghin et al., 2022). The amplitude of the P2 was higher for incongruent compared to congruent trials.

**Figure 4.**
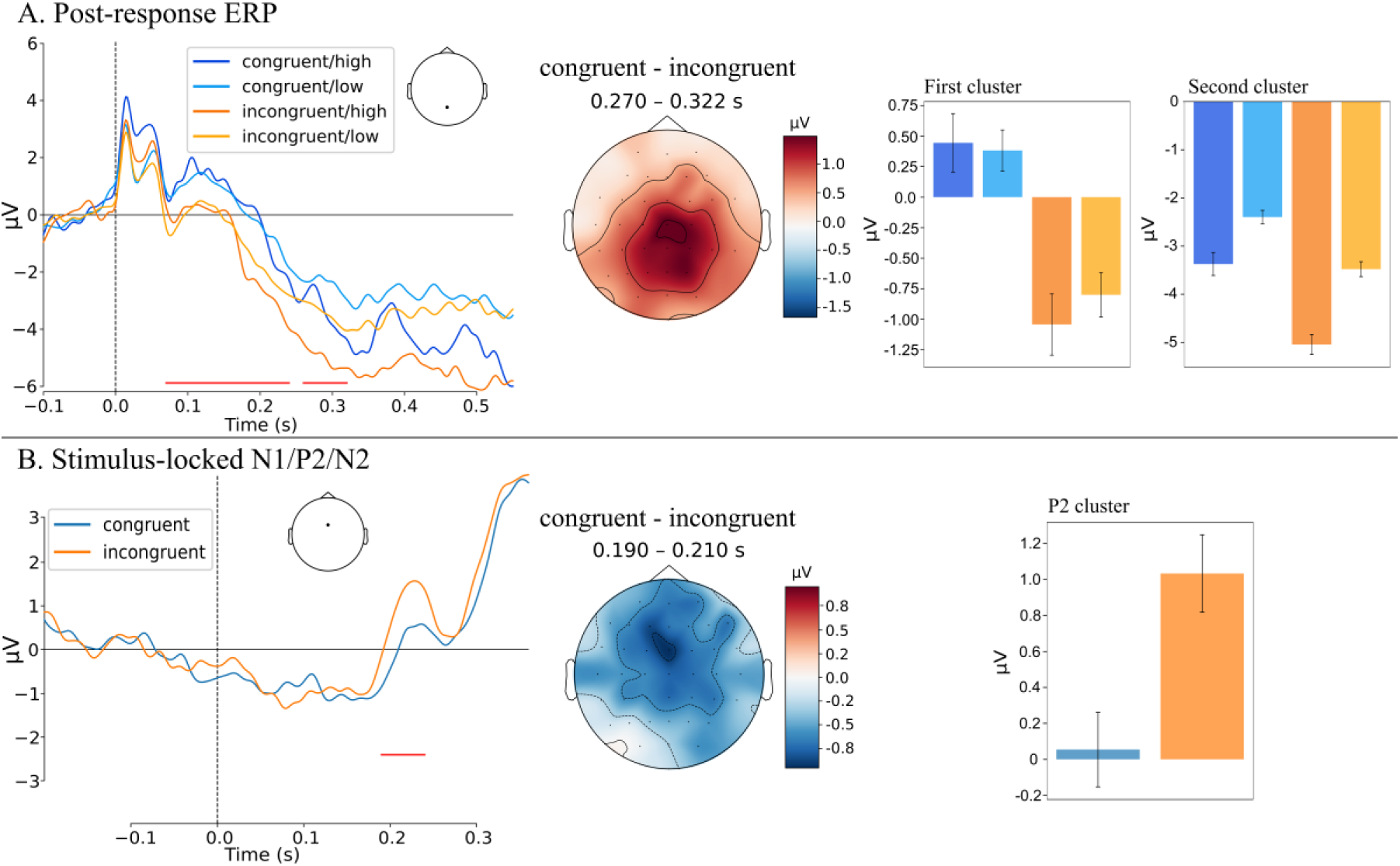
(**A, Left**) Grand average post-response ERPs at Pz, time-locked to the onset of the perceptual response (vertical dotted line), for congruent (orange lines) and incongruent (blue lines) trials, with low confidence (levels 1 and 2) shown in light colors and high confidence (levels 3 and 4) in dark colors. (**B, Left**) The graph depicts stimulus-locked ERP at FCz for the congruent (in orange) and incongruent (in blue) trials. Red horizontal lines in both graphs, represent significant time windows obtained with a cluster-based permutation test comparing congruent and incongruent trials. The topographies presented on the right panel depict the voltage difference between congruent and incongruent trials averaged across the time window of the temporal cluster observed in the response-locked (**A, Middle**) and the stimulus-locked (**B, Middle**) ERPs. Note that for the response-locked ERPs, the second cluster is depicted. The bar plots in the right panel depict the across-participant average amplitude of post-response activity for the two temporal clusters (**A, Right**) and post-stimulus activity at P2 (**B, Right**). Vertical bars represent the standard error of the mean.

Since there was a small effect of congruency on accuracy in Experiment 3, we performed the same tests on correct trials only, which yielded to similar results (see SI, *Figure S7*). We observed no modulation of congruency on the N2. Similarly, post-stimulus theta oscillations did not vary between spatially congruent and spatially incongruent trials (see SI for a discussion of these null results).

**Post-response activity.** Further analyses investigated the influence of spatial congruency on post-response signals. Firstly, we conducted a cluster-based permutation test comparing congruent and incongruent EEG epochs going from 0 to 500 ms post-response at Pz (other electrodes and group of electrodes were used for these analyses, all leading to similar results, see SI). This analysis revealed two significant temporal clusters (Figure 2A), one spanning from 70 ms to 240 ms, and another one from 260 ms to 320 ms. Subsequently, we categorized trials into low confidence (levels 1 and 2) and high confidence (levels 3 and 4) trials. To investigate interactions between confidence and congruency, we conducted a 2 (congruency: congruent vs. incongruent) by 2 (confidence: high vs. low) repeated-measures ANOVA on the average amplitude of each temporal cluster. The analyses of the early cluster (i.e., from 70 to 240 ms) showed a main effect of congruency (F(23) = 10.66, p = 0.003, Figure 2A), with a more negative activity for incongruent compared to congruent trials. Neither the main effect of confidence nor the interaction was significant (F(23) = 0.03, p = 0.86 and F(23) = 0.25, p = 0.62, respectively). In the second temporal cluster (i.e., from 260 to 320 ms) we observed a main effect of both congruency (F(23) = 8.71, p = 0.007) and confidence (F(23) = 5.05, p = 0.03, Figure 2A). The EEG activity was more negative for incongruent compared to congruent, and for high compared to low confidence trials. No significant congruency and confidence interaction were observed (F(23) = 0.37, p = 0.55). Since there was a small behavioral effect of congruency on accuracy, we performed the same EEG analyses on correct trials only and we observed similar results (see SI, Figure S7).

**LRP results.** The jackknife-based analysis revealed that the congruent LRP started slightly earlier (interpolated estimate = −184 ms) compared to the incongruent LRP (interpolated estimate = −157 ms), showing a difference in onset latency of 27 ms (SD = 7.2, t(23) = 3.77, p < 0.01, Figure 5). These findings, along with the observed behavioral effect of congruency on response times (cf. Experiments 2 and 3), suggest that the motor priming task successfully induced action preparation prior to Gabor presentation.

**Figure 5.**
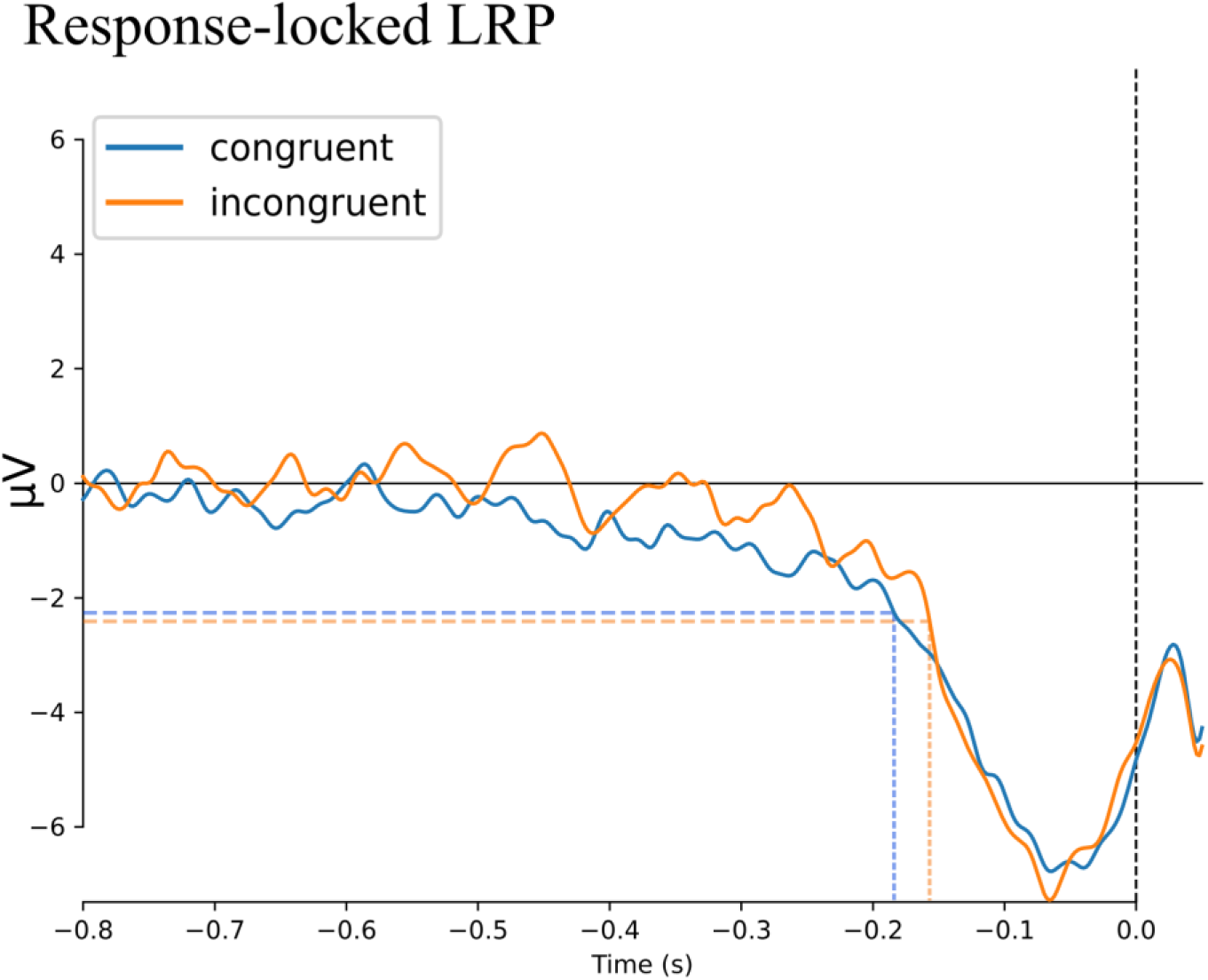
Response-locked Lateralized Readiness Potential for the congruent (red line) and incongruent (blue line) trials prior to the onset of the perceptual response (vertical dotted black line). The colored dotted lines represent the LRP onset latency at 1/3 threshold of the peak amplitude for congruent (blue) and incongruent (red) trials extracted from a Jackknife-based procedure showing a significant difference of 27 ms in LRP onset latencies.

## Discussion

Motor processes engaged during perceptual decisions have been found to affect retrospective confidence judgments. In three pre-registered experiments, we tested two hypotheses regarding how motor preparation impacts perceptual confidence. The “fluency hypothesis” suggests that the ease and smoothness of selecting and executing a motor response may serve as a cue for confidence judgments (Fleming et al., 2015). In contrast, the “monitoring hypothesis” proposes that confidence is influenced by action monitoring processes; it is not the fluency of the action, but rather the degree of control and the level of monitoring required for implementing a perceptual decision that affect confidence (cf. Sanchez et al., 2024). To test these hypotheses, participants reported the orientation of a Gabor stimulus (vertical or horizontal) and judged their confidence in their decision. Using a novel motor priming paradigm, participants were prompted to prepare an action in advance, which could be either compatible or incompatible with the action ultimately chosen to report the Gabor orientation.

In the three experiments, spatial congruency consistently influenced confidence judgments. Specifically, trial-by-trial analyses with mixed-effect models controlling for accuracy and response time revealed that participants were more confident in their decision when their response was spatially incongruent with the primed action (see Figure 3 and Table 1). This effect was general and independent of whether we primed the same (index fingers) or a different effector (feet in Experiment 1, and middle fingers in Experiment 2). Accordingly, these results suggest that early action preparation, involving spatial representation but not effector-specific representation, significantly influenced perceptual confidence. The fact that spatial representation of motor planning impacted response time and confidence can be explained by the partial overlap of spatial and semantic codes across different effectors and different actions (Fournier et al., 2010). Additional evidence that action preparation was modulated specifically by spatial congruence, rather than by effector compatibility, comes from the fact that responses were faster in spatially congruent compared to spatially incongruent trials (cf. Experiments 2 and 3). Furthermore, we observed an earlier beginning of the Lateralized Readiness Potentials (LRP) in spatially congruent compared to spatially incongruent responses. LRP activity has been typically attributed to motor preparation activity (Schmitz et al., 2019). Accordingly, these findings may suggest that the faster response times for congruent responses result from earlier motor preparation, whereas incongruent responses may involve delayed motor preparation (cf. Mordkoff & Gianaros, 2000). Overall, the findings on response times and LRP, confirms that our priming paradigm successfully influenced the selection/preparation of perceptual response, particularly for spatially congruent actions.

At first glance, the confidence results might seem compatible with many theories linking confidence to the accumulation of evidence for a decision (see Desender et al., 2021, for a recent investigation of confidence within evidence accumulation models). The fact that participants were more accurate (in Experiment 3) and slower (in Experiments 2 and 3) for incongruent trials might suggest that they were more cautious in these trials (cf. Vickers & Packer, 1982), and that they were accumulating more evidence before making their decision, which would result in a higher confidence. However, this explanation is unlikely for two main reasons. First, longer response times for incongruent trials are more likely due to inhibitory processes at the motor planning level (as revealed by the LRP observed in these trials in Experiment 3) than to prolonged evidence accumulation. Second, and more importantly, longer evidence accumulation would predict an increase of confidence for correct responses but a decrease in confidence for errors. Instead, we observed a boost in confidence for both correct and incorrect incongruent trials, with this effect being stronger for errors than for correct trials in Experiment 1.

The observed confidence boost in incongruent trials provides stronger support for the monitoring hypothesis rather than the fluency hypothesis. In fact, the fluency hypothesis would have predicted the opposite result: that actions whose selection is facilitated (i.e., congruent actions) lead to higher confidence judgments than incongruent actions (cf. Fleming et al., 2015). Our findings align with our previous study showing that, even when controlling for response time, perceptual responses preceded by partial motor activations (either ipsilateral or contralateral) consistently yielded higher confidence (Gajdos et al., 2019). It has been shown that partial activations are followed by an Error-Related Negativity (Meckler et al., 2017) originating in the Supplementary Motor Area (Bonini et al., 2014), a structure involved in action monitoring (Coull et al., 2016). Hence, these results suggested that partial motor activations may reflect premature commitments to a response that are effectively controlled and inhibited, thereby resulting in a greater sense of control over one’s responses and thus boosting confidence.

Similarly, in our study, perceptual responses that were incongruent with planned actions may have been interpreted by the system as instances in which an unwanted motor plan was successfully controlled, thereby leading to an increased sense of control and confidence. In other words, the ability to successfully resolve response competition by engaging monitoring mechanisms (Desender et al., 2014; Questienne et al., 2018) informs confidence estimations.

Stimulus-locked ERPs observed in Experiment 3 provide further support for this explanation. Our analyses revealed higher amplitude of the P2 component in incongruent trials compared to congruent trials. The frontal P2 component is thought to reflect early attentional resource allocation (Xie et al., 2020) associated with cognitive control (Ghin et al., 2022), sensitive to early updating processes during task-switching (Capizzi et al., 2015), and related to inhibitory processing, with a higher P2 for successful inhibited actions (Senderecka et al., 2012). We argue that preparing incongruent actions resulted in a greater involvement of early attentional resources necessary for response control, thereby impacting confidence. Interestingly, a recent study showed that TMS pulses to the dorsolateral prefrontal cortex (dlPFC) at 200 ms after stimulus onset influence confidence, suggesting confidence computations begin before the perceptual decision is fully formed, challenging post-response models where confidence is computed only after a response (Xue et al., 2023). Our study corroborates this notion by suggesting that post-stimulus processes as early as 200 ms correlate with retrospective confidence.

Another interesting aspect of the modulation of spatial congruency on confidence is that this effect was also observed in Experiment 1, despite the motor priming task not directly influencing motor execution—i.e., no effect of spatial congruency on response times was observed. The absence of a congruency effect on response times can be explained by the fact that, in the perceptual task of Experiment 1, a noise patch was presented 800 ms before the Gabor stimulus, signaling to participants that the Gabor presentation was imminent. As a result, even though the speeded RT task may have initially prepared the system for a specific action, the 800 ms delay likely eliminated the impact of motor priming on motor execution. This is consistent with the well-established finding that priming effects decay over time (Van den Bussche et al., 2009). However, despite the lack of an observable effect on motor execution, the conflicting motor representation induced by the speeded RT task still modulated retrospective confidence. This finding suggests that the motor signal used by the system to modulate confidence operates relatively early in the motor processing hierarchy—specifically, at the level of early motor planning rather than late motor preparation or motor execution. Interestingly, Fleming et al. (2015) also found that early motor processes play a crucial role in confidence modulation, rather than late motor processes or motor execution itself.

Although the current study supports the monitoring hypothesis, the specific feature that triggers monitoring processes has yet to be identified. Two distinct, but not mutually exclusive, factors may be at play. One possibility is that the effect stems from conflicting motor plans. Alternatively, the conflict driving monitoring and changes in confidence may arise between a motor plan and the perceptual decision the participant intends to commit to. Further research is needed to address this issue. While the exact feature triggering monitoring and affecting confidence is still unclear, our results more generally align with several studies demonstrating that cues and heuristics, such as stimulus visibility cue (Shekhar and Rahnev, 2024), response time (Kiani et al., 2014) or interoceptive states (Allen et al, 2016) do contribute to people’s sense of confidence beyond the strength of accumulated evidence.

Our findings are in apparent contrast with recent research supporting the fluency hypothesis. Fleming et al. (2015) showed that TMS stimulation over premotor areas linked to the chosen perceptual response enhances confidence, whereas incongruent stimulation decreases confidence. A critical difference between Fleming et al.’s study and ours may lie in the presence of response conflict. In their study, stimulation of the premotor area may have facilitated response planning without inducing implicit or explicit conflict with the action selected to report the stimulus. Participants may not have even perceived any difference between priming a congruent or incongruent action. In contrast, our study involved tasks with a certain degree of action conflict, thereby more strongly engaging monitoring and action control processes.

It could be argued that both Fleming et al.’s study and ours rely, after all, on similar processes, such as individuals’ control over their actions. Research on action fluency has shown that subliminal motor priming reduces participants’ sense of control, when the prime is incongruent with the performed action, but it increases it when the prime is congruent (Sidarus, Chambon, & Haggard, 2013; Chambon & Haggard, 2012; Wenke, Fleming, & Haggard, 2010). A similar interpretation may apply to Fleming et al.’s (2015) study, where participants may have experienced more control over fluent (congruent) responses compared to incongruent TMS stimulations. Interestingly, recent findings suggest that tasks involving response conflict, and thus requiring more cognitive control, increase individuals’ sense of control over their actions (Van den Bussche et al., 2020). Therefore, one might argue that in our experiment, a greater sense of control occurred during incongruent actions, as participants successfully resolved a conflict by controlling and inhibiting unwanted responses. In essence, the increase in perceptual confidence observed both in our study and in Fleming et al.’s study may rely on an increased sense of control over one’s actions. However, these experiences of control are triggered by distinct signals. Further studies should explore the relationship between sense of control and perceptual confidence.

Converging results regarding the influence of spatial congruency on confidence were also observed when examining post-response brain activity. Specifically, spatial congruency appeared to modulate post-decisional markers of response evaluation in two separate time windows. The first time-window spanned from 70 ms to 240 ms after the perceptual response. Activity within this period typically reflects early response monitoring processes, such as those underlying the Error-Related Negativity (ERN; Scheffers & Coles, 2000), a waveform observed following the detection of an incorrect choice. In the present study, this first post-response temporal window was solely modulated by congruency, and not by confidence. We argue that activity within this period reflects an early retrospective evaluation of the motor response, signaling the resolution of a motor conflict and the successful control of an unwanted motor plan. It is important to note that we could not identify a typical ERN component in our study (see SI). The ERN is classically observed in tasks (e.g., Simon task, Flankers task, etc.) where errors are explicitly detected by the participants. However, instances where participants realized they made a mistake were very rare in our experiment (see Methods). Therefore, we cannot and do not attempt to attribute our effect specifically to the ERN.

The second time window spanned from 260 ms to 320 ms after the response, where the Error Positivity (Pe; Pereira et al., 2020) is typically observed. The Pe is an event-related component characterized by a positive amplitude following incorrect choices. It is regarded as a reliable indicator of an individual’s ability to monitor errors during decision-making tasks (Davies et al., 2001). Recent studies have identified a family of ERP waves within the time window traditionally associated with the Pe. These waves not only reflect error monitoring but also appear to encode different levels of confidence (Boldt & Yeung, 2015). Specifically, within the time window where the classical Pe is observed, there are graded changes in voltage amplitude, corresponding to varying levels of confidence. Higher confidence levels are associated with more negative amplitudes, whereas lower levels of confidence and the certainty of error tend to manifest as more positive values.

In line with these observations, the second post-response temporal cluster we observed covaried not only with spatial congruency but also with confidence, consistent with previous results on the Pe (Boldt & Yeung, 2015; see SI). Specifically, high-confidence judgments were associated with more negative amplitudes compared to low-confidence trials. Hence, we argue that this second temporal cluster reflects a metacognitive evaluation of response accuracy. We would like to highlight that as for the ERN, we did not identify a typical Pe, since trials in which participants acknowledged they made an error were very rare.

Surprisingly, preparing the same or a different effector than the one used to report the Gabor did not seem to impact response times and confidence. We believe that the absence of this effect can be explained by the experimental design. For example, in the first experiment, we observed a clear cost when preparing foot actions and reporting the Gabor with the hands. Notably, accuracy decreased in this condition compared to when both the speeded RT task and the perceptual task required hand responses. If incongruent motor plans tend to increase confidence (because they require more monitoring), while incorrect responses reduce confidence, then these two effects may have cancelled each other out. Actually, in the different-effector block of the first experiment, not only did perceptual accuracy decrease, but metacognitive accuracy also declined. This suggests that the cost of switching from foot to hand responses impaired both perceptual and metacognitive evaluations, as if participants were distracted by the transition from lower to upper limb responses. In the second experiment, we aimed to eliminate the cost associated with feet responses by replacing these effectors with the middle fingers. Similarly, we observed no modulation of effector-specific representations on confidence or reaction times, but we observed a clear effect of spatial congruency on these two variables. This may be due to the stronger overlapping of motor codes for actions performed with the same hand than for actions performed with two different hands. In other words, for example, switching from left to right hand responses would likely generate more conflict than switching from the index to the middle finger of the same hand. Further studies may address these difficulties by employing, for example, hand actions and eye movements.

In summary, our results indicate that motor processes contribute, beyond perceptual evidence, in shaping retrospective perceptual confidence. Action planning modulate confidence through the involvement of monitoring systems (cf. also Yeung & Summerfield 2012). According to this framework, the system continuously monitors sensory and action information, with confidence estimates influenced by how effectively the system inhibits and controls alternative and unwanted responses (Gajdos et al., 2019; Anzulewicz et al., 2019). Thus, action processes are not merely ancillary processes for decision implementation but actively contribute to the formation of high-level cognitive constructs such as retrospective confidence.

## Constraints on Generality

The participants in this study were predominantly neurotypical young adults, a demographic often associated with greater cognitive flexibility and relatively high levels of digital literacy. Given the nature of recruitment methods, which were likely to attract educated individuals, these findings may not fully generalize to populations with different neurological profiles, age ranges, or educational backgrounds. Consequently, caution is warranted in extending these results to older adults, neurodivergent individuals, or those from less digitally engaged or lower educational backgrounds.

## Author Note

Conflicts of interest: all authors declare no conflicts of interest. Funding: this research was funded by the Agence Nationale de la Recherche, research funding ANR JCJC, project number ANR-18-CE10-0001. Artificial intelligence: No artificial intelligence assisted technologies were used in this research or the creation of this article. Ethics: this research complies with the Declaration of Helsinki (2023) and approved by the ethics committee of Université Paris Descartes (authorization n. 00012023-101). We are grateful to Dr Emmanuelle Bonnet for her help in the acquisition of the EEG data, and to Dr Louise Barne for her help in the pre-processing of EEG signals. We would also like to thank Profs. Steve Fleming, Nick Yeung and Mathieu Servant for their comments and suggestions on this research.

Ideas and results were presented at several conferences: 1) Investigating the role of motor preparation in perceptual confidence, *GDR Vision 2021*, Lilles, France; 2) Action planning modulates perceptual confidence but not perceptual accuracy, *Cognitive Neuroscience Society Annual Meeting 2022*, San Francisco, USA; 3) Action planning modulates perceptual confidence, *International Conference of Cognitive Neuroscience 2022*, Helsinki, Finland; 4) Action monitoring contributes to perceptual confidence: insights from EEG signals, *Association for the Scientific Study of Consciousness 2022*, Amsterdam, Netherlands.

