## Supplementary Information for "Action planning modulates perceptual confidence through action monitoring processes"

### Experimental procedure

**Participant exclusion.** No data were analyzed before completion of the experiment, except using a R script to verify the data validity based on our exclusion criteria. The protocols, data exclusion procedure and the R scripts were pre-registered on Open Science Framework before starting the experiments (https://osf.io/z6v9y). We expected participants to use the four levels of confidence ratings, so we excluded participants that did not use all confidence levels at least once. Based on this criterion, one participant was excluded in Experiment 2 (s/he used the confidence level 4). In Experiment 1, the task was interrupted for three participants during the training phase because they reported not being able to perform the double task, as the required speeded response timing was too fast for them. Following the preregistered exclusion procedure, 10 participants were excluded in Experiment 1 because their perceptual performance was either too high (3 participants above 90% accuracy) or too low (7 participants under 55% accuracy). For the same reason, 8 participants were excluded in Experiment 2, all of them had an average perceptual accuracy below 60%. No participants were excluded in Experiment 3. Importantly, we replicated the analyses while including all the participants excluded in Experiment 1 and 2 and the statistical results were qualitatively identical (see Table S6, S7 & S8).

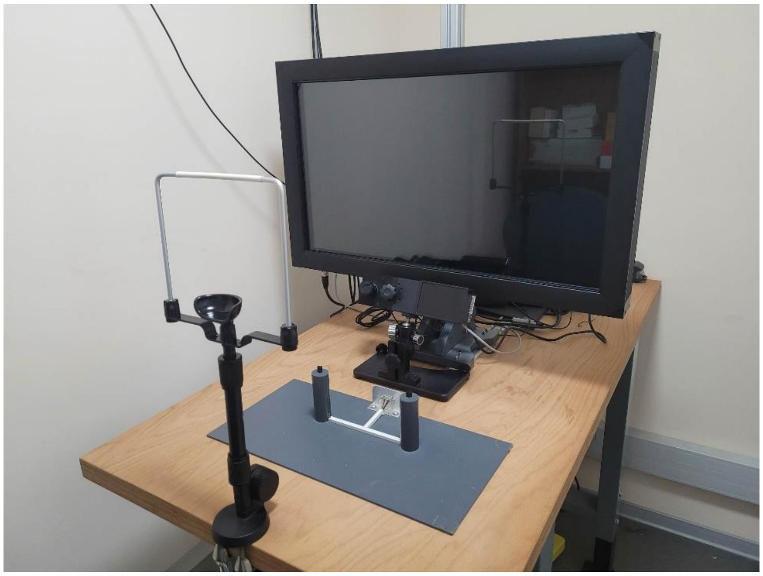

**Figure S1**. Experiment 3 set up and button-box for thumb responses.

**Preliminary calibration phase.**
Before starting the main experiment, we determined participants' orientation discrimination thresholds using a triple interleaved staircase. This method allowed us to identify Gabor contrast values associated with a 75% discrimination accuracy, a level known to be appropriate for studying confidence (Ferrigno, Bueno, & Cantlon, 2019; Fleming et al., 2015). Each staircase comprised 40 trials, employing an adaptive procedure known as accelerated stochastic approximation (Kesten, 1958). In staircase trials, participants were not required to report confidence judgments.
Prior to the staircase, participants completed 60 staircase training trials in which they were familiarized with the stimuli and the perceptual task. After this training, participants performed 120 staircase trials (i.e., 3 interleaved staircases x 40 trials). Subsequently, they were familiarized to the primary task with 30 training trials.
Calibration aimed for 70–80% accuracy. Due to a high failure rate in the first experiment (38% of participants), a supplementary phase was added in the second and third experiments. After psychometric staircases, the contrast value was tested on 20 perceptual trials with confidence ratings, repeating until validation criteria were met: 1. accuracy between 60–90% (2–8 errors), 2. no more than six "very confident" (level 4) confidence ratings to prevent excessive visibility, and 3. no more than six "random guess" (level 1) confidence ratings to prevent invisibility. If criteria were unmet, contrast was adjusted by 0.0025, and trials repeated.

### Data analyses and results

**Mixed models.** Data were analyzed in R (R Core Team 2021) using mixed-effects models available in the lme4 package (Bates et al., 2015). Response times (RT) were calculated with respect to the onset of the Gabor patch. The tables below (S1 and S2) report in full the results relative to response times and accuracy observed in the three experiments.

**Description and discussion of interactions modulating confidence.**
In Experiment 1, we found an interaction between effector compatibility and accuracy (odds ratio = 1.08, $p < 0.001$, see SI, Figure S2 & Post-hoc tests), indicating poorer metacognitive accuracy in the ‘different effector’ block (using foot responses) compared to the ‘same effector’ block. Participants were more confident about incorrect responses in the ‘different effector’ block (M = 1.71, SE = 0.14) than in the ‘same effector’ block (M = 1.60, SE = 0.11), and less confident when correct in the ‘different effector’ block (M = 2.53, SE = 0.14) than in the ‘same effector’ block (M = 2.71, SE = 0.13). A possible explanation is that switching from feet to hand responses was more challenging and costly compared to keep responding with the hands, thus deteriorating both perceptual accuracy and metacognitive accuracy. This interaction disappeared in Experiment 2, where we replaced foot responses with middle finger responses, thereby removing the potential cost of switching from lower to upper limbs in the different effector block. Secondly, effector compatibility interacted with effector order in Experiment 1 (odds ratio = 0.88, $p = 0.003$, see SI, Figure S2 & Post-hoc tests), showing that confidence decreased for the ‘different effector’ block when participants began with the ‘same effector’ block. We speculate that this modulation of confidence might be due to fatigue, combined with the cost of shifting from upper to lower limb responses, since the task is more difficult for the ‘different effector’ block. Accordingly, increased fatigue may reduce overall confidence (see also main effect of effector order on confidence), particularly when participants must also divide attention between lower and upper limbs. Note that we found no evidence for an effect in Experiment 2 where we removed feet responses.

Finally, only in Experiment 1 we observed a three-way interaction between effector order, effector and accuracy (odds ratio = 1.08, $p < 0.001$, see SI, Figure S2 & Post-hoc tests). Overall, confidence decreased in the “different effector” block compared to the “same effector” block, except when participants completed the “different effector” block first and made incorrect perceptual decisions. In this specific case, the “different effector” block elicited higher confidence than the “same effector” block. We do not have a clear explanation for this finding. As with other interactions involving effector compatibility, this interaction was not observed in Experiment 2.

**Table S1.** Hierarchical regression tables predicting **accuracy** from congruency and reaction time for Experiment 3 (right), plus effector and effector order for Experiment 1 (left), 2 (middle), and 3 (right). Coefficients were coded as follows: Accuracy: error = -1, correct = 1; Congruency: incongruent = -1, congruent = 1; Effector compatibility: different effector (feet in Experiment 1, middle fingers in Experiment 2) = -1, same effector (index fingers) = 1; Bloc order: same effector first = -1, different effector first = 1. Response time is mean-centered. The precise models are the following logit mixed-effect models:
Experiment 1: accuracy ~ congruency * effector * effector_order * rt + (1 + rt || subject)
Experiment 2: accuracy ~ congruency * effector * effector_order * rt + (1 + effector + rt || subject)
Experiment 3: accuracy ~ congruency * rt + (1 + rt || subject)

| | **Accuracy (Exp. 1)** | | | **Accuracy (Exp. 2)** | | | **Accuracy (Exp. 3)** | | |
|---|---|---|---|---|---|---|---|---|---|
| *Predictors* | *Odds Ratios* | *CI* | *p* | *Odds Ratios* | *CI* | *p* | *Odds Ratios* | *CI* | *p* |
| (Intercept) | 2.22 | 1.76 – 2.80 | **<0.001** | 3.06 | 2.57 – 3.64 | **<0.001** | 2.92 | 2.45 – 3.47 | **<0.001** |
| congruency1 | 1.00 | 0.93 – 1.07 | 0.905 | 0.94 | 0.88 – 1.01 | 0.096 | 0.92 | 0.86 – 0.99 | **0.021** |
| effector1 | 1.13 | 1.06 – 1.21 | **<0.001** | 0.96 | 0.81 – 1.15 | 0.690 | | | |
| effector order1 | 1.04 | 0.82 – 1.31 | 0.769 | 0.92 | 0.77 – 1.10 | 0.349 | | | |
| rt gabor centered | 1.97 | 1.01 – 3.82 | **0.046** | 0.19 | 0.10 – 0.37 | **<0.001** | 0.23 | 0.14 – 0.39 | **<0.001** |
| congruency1 * effector1 | 1.01 | 0.94 – 1.08 | 0.812 | 1.00 | 0.93 – 1.07 | 0.923 | | | |
| congruency1 * effector order1 | 1.03 | 0.96 – 1.11 | 0.361 | 1.00 | 0.93 – 1.07 | 0.968 | | | |
| effector1 * effector order1 | 1.06 | 0.99 – 1.14 | 0.089 | 1.12 | 0.93 – 1.33 | 0.227 | | | |
| congruency1 * rt gabor centered | 0.80 | 0.47 – 1.36 | 0.409 | 0.97 | 0.66 – 1.42 | 0.877 | 1.74 | 1.26 – 2.39 | **0.001** |
| effector1 * rt gabor centered | 0.96 | 0.56 – 1.64 | 0.887 | 0.95 | 0.62 – 1.47 | 0.825 | | | |
| effector order1 * rt gabor centered | 0.36 | 0.18 – 0.70 | **0.003** | 0.88 | 0.45 – 1.71 | 0.704 | | | |
| congruency1 * effector1 * effector order1 | 0.99 | 0.92 – 1.06 | 0.746 | 0.97 | 0.91 – 1.04 | 0.450 | | | |
| (congruency1 * effector1) * rt gabor centered | 1.15 | 0.67 – 1.95 | 0.613 | 1.39 | 0.95 – 2.03 | 0.086 | | | |
| (congruency1 * effector order1) * rt gabor centered | 0.95 | 0.56 – 1.61 | 0.848 | 1.13 | 0.78 – 1.66 | 0.512 | | | |
| (effector1 * effector order1) * rt gabor centered | 0.94 | 0.55 – 1.61 | 0.834 | 0.84 | 0.54 – 1.29 | 0.424 | | | |
| (congruency1 * effector1 * effector order1) * rt gabor centered | 1.02 | 0.60 – 1.74 | 0.935 | 1.09 | 0.75 – 1.59 | 0.647 | | | |

**Table S2.** Hierarchical regression tables predicting **reaction times** from accuracy and congruency for Experiment 3 (right), plus effector and effector order for Experiment 1 (left), 2 (middle), and 3 (right). Coefficients were coded as follows: Accuracy: error = -1, correct = 1; Congruency: incongruent = -1, congruent = 1; Effector compatibility: different effector (feet in Experiment 1, middle fingers in Experiment 2) = -1, same effector (index fingers) = 1; Bloc order: same effector first = -1, different effector first = 1. Response time is mean-centered. . The precise models are the following generalized inverse-gaussian mixed-effect models (with identity link function):
Experiment 1: rt ~ accuracy * congruency * effector * effector_order + (1 + accuracy + effector || subject)
Experiment 2: rt ~ accuracy * congruency * effector * effector_order + (1 + accuracy + effector + effector_order || subject)
Experiment 3: rt ~ accuracy * congruency + (1 + accuracy_gabor + congruency || subject)

| | rt gabor*1000 | | | rt gabor*1000 | | | rt gabor*1000 | | |
|---|---|---|---|---|---|---|---|---|---|
| *Predictors* | *Estimates* | *CI* | *p* | *Estimates* | *CI* | *p* | *Estimates* | *CI* | *p* |
| (Intercept) | 470.66 | 422.18 – 519.13 | **<0.001** | 734.09 | 669.56 – 798.62 | **<0.001** | 732.44 | 691.49 – 773.39 | **<0.001** |
| accuracy gabor [1] | 6.69 | -2.50 – 15.88 | 0.153 | -18.16 | -23.48 – -12.84 | **<0.001** | -19.51 | -31.62 – -7.39 | **0.002** |
| congruency1 | -2.83 | -10.23 – 4.56 | 0.453 | -11.51 | -16.84 – -6.19 | **<0.001** | -13.69 | -23.80 – -3.59 | **0.008** |
| effector1 | 4.23 | -7.19 – 15.65 | 0.468 | -5.72 | -11.05 – -0.39 | **0.036** | | | |
| effector order1 | 7.67 | -24.07 – 39.41 | 0.636 | 14.68 | -38.14 – 67.50 | 0.586 | | | |
| accuracy gabor [1] × congruency1 | -0.25 | -7.62 – 7.12 | 0.947 | 0.83 | -4.45 – 6.11 | 0.759 | 7.86 | 1.93 – 13.78 | **0.009** |
| accuracy gabor [1] × effector1 | 0.52 | -7.05 – 8.09 | 0.893 | -1.87 | -7.20 – 3.45 | 0.491 | | | |
| congruency1 × effector1 | -0.67 | -8.26 – 6.91 | 0.862 | 2.02 | -3.25 – 7.29 | 0.453 | | | |
| accuracy gabor [1] × effector order1 | -8.51 | -17.64 – 0.62 | 0.068 | -0.98 | -6.31 – 4.35 | 0.717 | | | |
| congruency1 × effector order1 | -0.00 | -7.41 – 7.40 | 0.999 | -3.74 | -9.05 – 1.58 | 0.168 | | | |
| effector1 × effector order1 | -31.38 | -42.54 – -20.23 | **<0.001** | -0.94 | -6.29 – 4.42 | 0.732 | | | |
| accuracy gabor [1] × congruency1 × effector1 | 2.98 | -4.47 – 10.43 | 0.433 | 1.78 | -3.49 – 7.06 | 0.508 | | | |
| accuracy gabor [1] × congruency1 × effector order1 | 0.51 | -6.91 – 7.93 | 0.892 | 0.36 | -4.91 – 5.64 | 0.893 | | | |
| accuracy gabor [1] × effector1 × effector order1 | 0.91 | -6.74 – 8.56 | 0.815 | -1.01 | -6.36 – 4.34 | 0.711 | | | |
| congruency1 × effector1 × effector order1 | -0.34 | -7.84 – 7.17 | 0.930 | 4.47 | -0.83 – 9.77 | 0.098 | | | |
| accuracy gabor [1] × congruency1 × effector1 × effector order1 | -2.25 | -9.88 – 5.38 | 0.563 | -0.84 | -6.13 – 4.45 | 0.755 | | | |

**Post-hoc tests.** Post-hoc tests using the package emmeans were performed to explore all significant interactions. P-values were corrected according to the False Discovery Rate (FDR) method. The results of the comparisons are reported below.

**Experiment 1: interactions effects on confidence (see Figure S2).** Note that for interactions involving accuracy, we did not report paired comparisons between correct and incorrect responses, as correct responses consistently elicited higher confidence than incorrect ones. Therefore, for post-hoc tests, we report only comparisons where accuracy is held constant while varying the second factor in the interaction.

1. **Post-hoc investigating the interaction effect of accuracy and congruency on confidence**

```
**Accuracy is constant**

accuracy_gabor = 0:
 contrast               estimate     SE  df z.ratio p.value         p adjusted
 congruent - incongruent  -0.2132 0.0701 Inf -3.039  0.0024         0.0048

accuracy_gabor = 1:
 contrast               estimate     SE  df z.ratio p.value         p adjusted
 congruent - incongruent  -0.0381 0.0457 Inf -0.835  0.4039         0.4039

Results are averaged over the levels of: effector, effector_order
```

2. **Post-hoc investigating the interaction effect of accuracy and effector on confidence**

```
**Accuracy is constant**

accuracy_gabor = 0:
 contrast     estimate     SE  df z.ratio p.value p adjusted
 other - same    0.139 0.0969 Inf  1.433  0.1518  0.1518

accuracy_gabor = 1:
 contrast     estimate     SE  df z.ratio p.value p adjusted
 other - same   -0.183 0.0803 Inf -2.279  0.0227  0.0454

Results are averaged over the levels of: congruency, effector_order
```

3. **Post-hoc investigating the interaction effect of effector and effector order on confidence**

```
**Effector_order is constant**

effector_order = other1:
 contrast     estimate    SE  df z.ratio p.value  p adjusted
 other - same    0.224 0.114 Inf  1.971  0.0488   0.065

effector_order = same1:
 contrast     estimate    SE  df z.ratio p.value  p adjusted
 other - same   -0.268 0.114 Inf -2.345  0.0190   0.0380

Results are averaged over the levels of: accuracy_gabor, congruency


**Effector is constant**

effector = other:
 contrast       estimate    SE  df z.ratio p.value         p adjusted
 other1 - same1   -0.256 0.292 Inf -0.875  0.3816         0.3816

effector = same:
 contrast       estimate    SE  df z.ratio p.value         p adjusted
 other1 - same1   -0.748 0.226 Inf -3.314  0.0009         0.0036

Results are averaged over the levels of: accuracy_gabor, congruency
```

4. **Post-hoc investigating the interaction effect of effector, effector order and accuracy on confidence**

```
**Effector and Accuracy are constant**

effector = other, accuracy_gabor = 0:
 contrast       estimate    SE  df z.ratio p.value         p adjusted
 other1 - same1   0.0222 0.363 Inf  0.061  0.9512          0.9512

effector = same, accuracy_gabor = 0:
 contrast       estimate    SE  df z.ratio p.value         p adjusted
 other1 - same1  -0.7670 0.294 Inf -2.605  0.0092          0.0245

effector = other, accuracy_gabor = 1:
 contrast       estimate    SE  df z.ratio p.value         p adjusted
 other1 - same1  -0.5332 0.297 Inf -1.796  0.0725          0.0967

effector = same, accuracy_gabor = 1:
 contrast       estimate    SE  df z.ratio p.value         p adjusted
 other1 - same1  -0.7287 0.256 Inf -2.849  0.0044          0.0176

Results are averaged over the levels of: congruency


**Effector_order and Accuracy are constant**

effector_order = other1, accuracy_gabor = 0:
 contrast     estimate    SE  df z.ratio p.value  p adjusted
 other - same   0.5335 0.140 Inf  3.824  0.0001   0.0008

effector_order = same1, accuracy_gabor = 0:
 contrast     estimate    SE  df z.ratio p.value  p adjusted
 other - same  -0.2558 0.137 Inf -1.862  0.0627   0.0967

effector_order = other1, accuracy_gabor = 1:
 contrast     estimate    SE  df z.ratio p.value  p adjusted
 other - same  -0.0853 0.115 Inf -0.744  0.4571   0.5224

effector_order = same1, accuracy_gabor = 1:
 contrast     estimate    SE  df z.ratio p.value  p adjusted
 other - same  -0.2807 0.117 Inf -2.398  0.0165   0.033

Results are averaged over the levels of: congruency
```

**Experiment 1: interactions effects on response times (see Figure S4).** The following post-hoc investigate the interactions effect on response times observed in Experiment 1.

**Post-hoc investigating the interaction effect of effector and effector order on response times**

```
effector_order = feet first:
contrast      estimate   SE  df  z.ratio   p.value
feet - hands     54.3   16.8 Inf   3.225   0.0013

effector_order = hands first:
contrast      estimate   SE  df   z.ratio  p.value
feet - hands    -71.2   15.7 Inf   -4.534  <.0001

Results are averaged over the levels of: accuracy_gabor, congruency
```

**Experiment 2: interactions effects on confidence (Figure S3).** The following post-hoc investigate the interactions effect on confidence observed in Experiment 2. None of the paired comparisons elicited a significant p-value (after correction for multiple comparison).

1. **Post-hoc investigating the interaction effect of accuracy, congruency and effector order on confidence**

```
**Effector_order and Accuracy are constant**

effector_order = other1, accuracy_gabor = 0:
 contrast                estimate     SE  df z.ratio p.value       p adjusted
 congruent - incongruent   0.0523 0.1098 Inf  0.476  0.6340        0.634

effector_order = same1, accuracy_gabor = 0:
 contrast                estimate     SE  df z.ratio p.value       p adjusted
 congruent - incongruent  -0.3071 0.1142 Inf -2.689  0.0072        0.0564

effector_order = other1, accuracy_gabor = 1:
 contrast                estimate     SE  df z.ratio p.value       p adjusted
 congruent - incongruent  -0.1379 0.0760 Inf -1.814  0.0697        0.1823

effector_order = same1, accuracy_gabor = 1:
 contrast                estimate     SE  df z.ratio p.value       p adjusted
 congruent - incongruent  -0.1296 0.0791 Inf -1.639  0.1012        0.1823

Results are averaged over the levels of: effector

**Congruency and Accuracy are constant**

congruency = congruent, accuracy_gabor = 0:
 contrast       estimate    SE  df z.ratio p.value         p adjusted
 other1 - same1   -0.468 0.333 Inf -1.407  0.1595          0.1823

congruency = incongruent, accuracy_gabor = 0:
 contrast       estimate    SE  df z.ratio p.value         p adjusted
 other1 - same1   -0.828 0.337 Inf -2.455  0.0141          0.056

congruency = congruent, accuracy_gabor = 1:
 contrast       estimate    SE  df z.ratio p.value         p adjusted
 other1 - same1   -0.540 0.355 Inf -1.523  0.1279          0.1823

congruency = incongruent, accuracy_gabor = 1:
 contrast       estimate    SE  df z.ratio p.value         p adjusted
 other1 - same1   -0.532 0.361 Inf -1.473  0.1407 0.1823
```

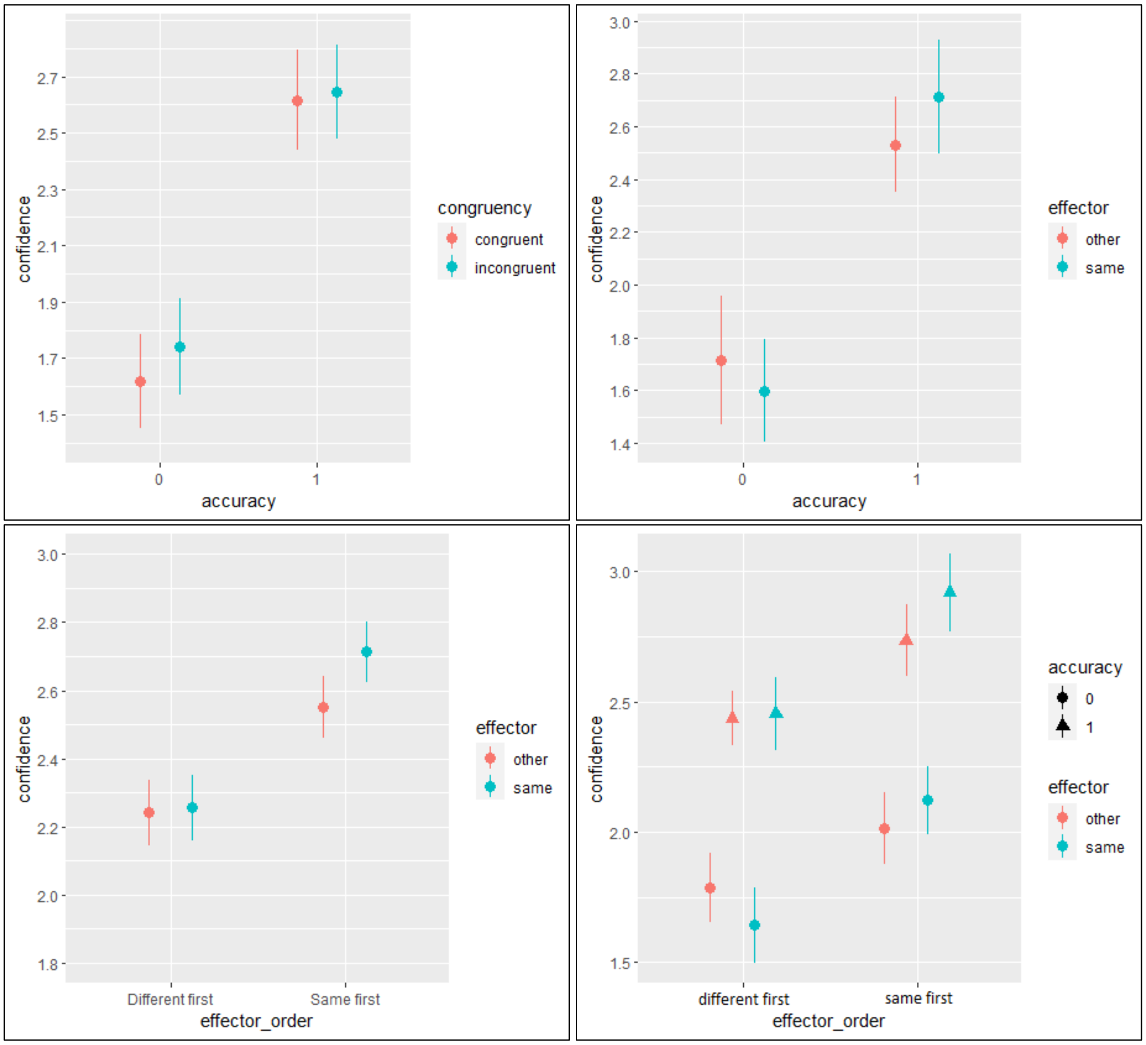


**Figure S2.** Confidence average values for Experiment 1 as a function of accuracy and congruency (top left); accuracy and effector (top right); effector and effector order (bottom left); effector order, accuracy, and effector (bottom right). Error bars represent confidence intervals indicating the dispersion of individual averages, corrected using the Cousineau & Morey method (Morey, 2008).

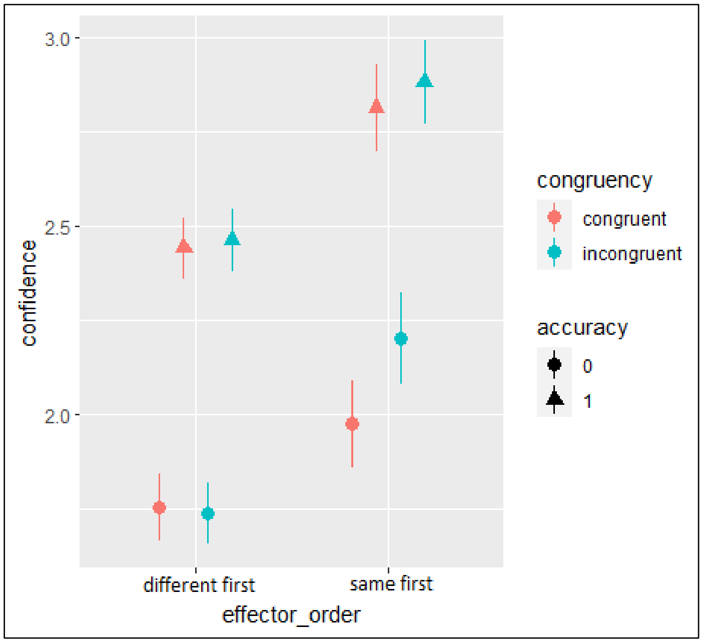


**Figure S3.** Confidence average values for Experiment 2 as a function of accuracy, effector order and congruency. Error bars represent confidence intervals indicating the dispersion of individual averages, corrected using the Cousineau & Morey method (Morey, 2008).

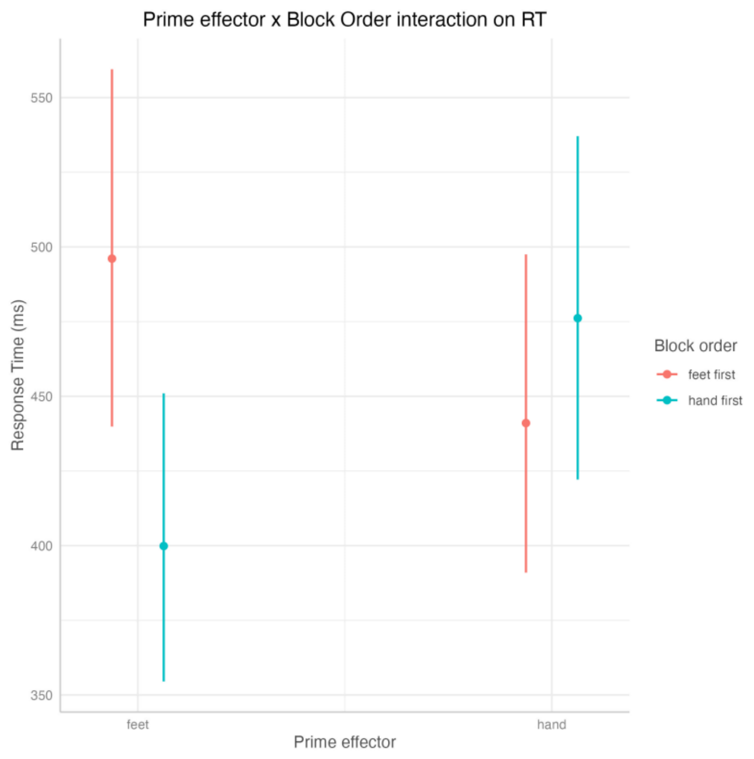


**Figure S4.** Interaction between effector and effector order from model predicting response times in Experiment 1. The error bars represent standard errors.

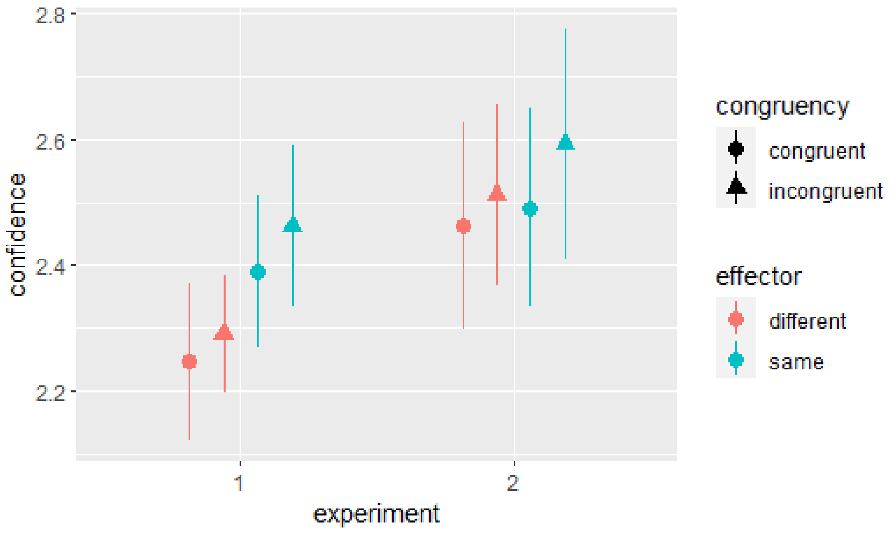


**Figure S5**. Confidence average values for Experiment 1 and Experiment 2 as a function of congruency and effector. Error bars represent confidence intervals indicating the dispersion of individual averages, corrected using the Cousineau & Morey method (Morey, 2008).

**Bayesian analyses of the behavior.** All analyses and results were replicated with Bayesian analyses implemented using the Brms package in R (Bürkner, 2017). Confidence was analyzed with ordered logistic models, accuracy was analyzed with Bernoulli models and RT with exgaussian models. In all cases, we used weakly informative priors for regression parameters.
Predictors were coded as follows: Accuracy: error = -1, correct = 1; Congruency: incongruent = -1, congruent = 1; Effector: different effector (feet or middle fingers for Experiment 1 and 2, respectively) = -1, same effector (hands or index fingers for Experiment 1 and 2, respectively) = 1; Effector Order: same effector first = -1, different effector first = 1. Reaction time is mean-centered.
Experiment 1: Number of subjects: 16. Number of observations: 4224.
Experiment 2: Number of subjects: 24. Number of observations: 4510.
Experiment 3: Number of subjects: 24. Number of observations: 4570.

| | **conf** | | **conf** | | **conf** | |
|---|---|---|---|---|---|---|
| *Predictors* | *Odds Ratios* | *CI (95%)* | *Odds Ratios* | *CI (95%)* | *Odds Ratios* | *CI (95%)* |
| Intercept[1] | 0.63 | 0.44 – 0.92 | 0.42 | 0.29 – 0.59 | 0.85 | 0.72 – 0.99 |
| Intercept[2] | 1.61 | 1.12 – 2.33 | 1.20 | 0.84 – 1.70 | 2.34 | 1.99 – 2.75 |
| Intercept[3] | 4.35 | 3.03 – 6.31 | 3.04 | 2.13 – 4.34 | 6.01 | 5.09 – 7.08 |
| accuracy gabor: accuracy gabor 1 | 1.90 | 1.60 – 2.26 | 1.42 | 1.31 – 1.54 | 1.58 | 1.41 – 1.76 |
| congruency1 | 0.94 | 0.90 – 0.98 | 0.93 | 0.88 – 0.99 | 0.89 | 0.85 – 0.94 |
| effector1 | 1.01 | 0.91 – 1.11 | 0.98 | 0.86 – 1.10 | | |
| effector_order1 | 0.84 | 0.59 – 1.21 | 0.77 | 0.54 – 1.10 | | |
| rt gabor centered | 0.68 | 0.31 – 1.46 | 0.09 | 0.05 – 0.16 | 0.15 | 0.10 – 0.25 |
| accuracy_gabor1:congruency1 | 1.04 | 1.01 – 1.09 | 1.00 | 0.96 – 1.04 | 1.03 | 0.99 – 1.07 |
| accuracy_gabor1:effector1 | 1.09 | 1.05 – 1.13 | 1.02 | 0.98 – 1.06 | | |
| congruency1:effector1 | 0.98 | 0.94 – 1.02 | 0.98 | 0.94 – 1.02 | | |
| accuracy_gabor1:effector_order1 | 0.93 | 0.79 – 1.11 | 1.03 | 0.94 – 1.11 | | |
| congruency1:effector_order1 | 1.04 | 0.99 – 1.08 | 1.04 | 0.98 – 1.10 | | |
| effector1:effector_order1 | 0.88 | 0.80 – 0.97 | 0.91 | 0.81 – 1.03 | | |
| accuracy_gabor1:congruency1:effector1 | 1.02 | 0.98 – 1.06 | 1.03 | 1.00 – 1.08 | | |
| accuracy_gabor1:congruency1:effector_order1 | 1.00 | 0.96 – 1.04 | 0.95 | 0.92 – 0.99 | | |
| accuracy_gabor1:effector1:effector_order1 | 1.07 | 1.03 – 1.11 | 1.00 | 0.96 – 1.04 | | |
| congruency1:effector1:effector_order1 | 1.00 | 0.96 – 1.04 | 1.02 | 0.98 – 1.06 | | |
| accuracy_gabor1:congruency1:effector1:effector_order1 | 1.00 | 0.97 – 1.04 | 0.99 | 0.95 – 1.03 | | |
| **Random Effects** | | | | | | |
| $\sigma^2$ | 0.11 | | 0.19 | | 0.08 | |
| $\tau_{00}$ | 1.08 | | 1.06 | | 0.91 | |
| ICC | 0.09 | | 0.15 | | 0.08 | |
| N | 16 $_{subject\_id}$ | | 24 $_{subject\_id}$ | | 24 $_{subject\_id}$ | |
| Observations | 4224 | | 4510 | | 4570 | |
| Marginal $R^2$ / Conditional $R^2$ | 0.232 / 0.395 | | 0.214 / 0.468 | | 0.183 / 0.293 | |

**Table S3.** Bayesian analyses. Predicting confidence from accuracy, congruency, effector, effector order, and reaction time in Experiment 1 (left), Experiment 2 (middle), and Experiment 3 (right). All models are maximal models, i.e. they include all terms and their interactions as random variables.

| | acc gabor num | | acc gabor num | | acc num | |
|---|---|---|---|---|---|---|
| *Predictors* | *Odds Ratios* | *CI (95%)* | *Odds Ratios* | *CI (95%)* | *Odds Ratios* | *CI (95%)* |
| Intercept | 2.26 | 1.67 – 3.07 | 3.09 | 2.53 – 3.79 | 2.92 | 2.41 – 3.55 |
| congruency1 | 0.99 | 0.92 – 1.07 | 0.94 | 0.88 – 1.01 | 0.92 | 0.86 – 0.99 |
| effector1 | 1.14 | 1.00 – 1.30 | 0.97 | 0.79 – 1.18 | | |
| effector_order1 | 1.03 | 0.76 – 1.39 | 0.92 | 0.75 – 1.13 | | |
| rt gabor centered | 1.74 | 0.80 – 3.56 | 0.23 | 0.11 – 0.47 | 0.26 | 0.15 – 0.45 |
| congruency1:effector1 | 1.01 | 0.94 – 1.08 | 1.00 | 0.93 – 1.07 | | |
| congruency1:effector_order1 | 1.04 | 0.96 – 1.11 | 1.00 | 0.93 – 1.07 | | |
| effector1:effector_order1 | 1.06 | 0.92 – 1.20 | 1.11 | 0.91 – 1.37 | | |
| congruency1:rt_gabor_centered | 0.81 | 0.48 – 1.37 | 0.95 | 0.65 – 1.40 | 1.71 | 1.25 – 2.35 |
| effector1:rt_gabor_centered | 0.95 | 0.55 – 1.67 | 0.95 | 0.62 – 1.47 | | |
| effector_order1:rt_gabor_centered | 0.43 | 0.20 – 0.98 | 0.87 | 0.45 – 1.78 | | |
| congruency1:effector1:effector_order1 | 0.99 | 0.92 – 1.07 | 0.97 | 0.91 – 1.05 | | |
| congruency1:effector1:rt_gabor_centered | 1.16 | 0.70 – 1.97 | 1.38 | 0.95 – 2.03 | | |
| congruency1:effector_order1:rt_gabor_centered | 0.93 | 0.55 – 1.55 | 1.13 | 0.77 – 1.66 | | |
| effector1:effector_order1:rt_gabor_centered | 1.15 | 0.66 – 2.08 | 0.85 | 0.55 – 1.31 | | |
| congruency1:effector1:effector_order1:rt_gabor_centered | 1.01 | 0.60 – 1.69 | 1.10 | 0.75 – 1.61 | | |
| **Random Effects** | | | | | | |
| $\sigma^2$ | 0.00 | | 0.01 | | 0.00 | |
| $\tau_{00}$ | 0.22 | | 0.19 | | 0.19 | |
| ICC | 0.02 | | 0.03 | | 0.02 | |
| N | $16_{\text{subject\_id}}$ | | $24_{\text{subject\_id}}$ | | $24_{\text{subject\_id}}$ | |
| Observations | 4224 | | 4510 | | 4570 | |
| Marginal $R^2$ / Conditional $R^2$ | 0.019 / 0.064 | | 0.028 / 0.074 | | 0.018 / 0.046 | |

**Table S4.** Bayesian analyses. Predicting accuracy from congruency, effectors, effector order, and reaction time in Experiment 1 (left), Experiment 2 (middle), and Experiment 3 (right).

| | rt gabor*1000 | | rt gabor*1000 | | rt gabor*1000 | |
|---|---|---|---|---|---|---|
| *Predictors* | *Estimates* | *CI (95%)* | *Estimates* | *CI (95%)* | *Estimates* | *CI (95%)* |
| Intercept | 458.15 | 422.08 – 494.66 | 713.25 | 680.10 – 745.76 | 690.55 | 649.97 – 730.72 |
| accuracy gabor: accuracy gabor 1 | 6.48 | 0.13 – 12.67 | -18.06 | -26.83 – -9.42 | -17.19 | -25.54 – -9.00 |
| congruency1 | -2.52 | -7.18 – 2.05 | -9.64 | -15.56 – -3.61 | -18.96 | -27.12 – -11.01 |
| effector1 | 4.78 | -5.09 – 14.47 | -13.07 | -31.18 – 5.00 | | |
| effector_order1 | 14.16 | -21.54 – 52.07 | 14.06 | -20.45 – 48.55 | | |
| accuracy_gabor1:congruency1 | -0.59 | -4.29 – 3.10 | -1.36 | -5.52 – 2.72 | 7.97 | 3.02 – 13.02 |
| accuracy_gabor1:effector1 | 0.05 | -3.76 – 3.82 | -2.90 | -7.21 – 1.35 | | |
| congruency1:effector1 | -0.01 | -3.75 – 3.73 | -0.88 | -5.06 – 3.20 | | |
| accuracy_gabor1:effector_order1 | -6.80 | -13.26 – -0.44 | -6.74 | -15.38 – 1.93 | | |
| congruency1:effector_order1 | 1.18 | -3.40 – 6.01 | -6.71 | -12.65 – -0.89 | | |
| effector1:effector_order1 | -32.66 | -41.99 – -23.35 | -2.21 | -19.63 – 15.18 | | |
| accuracy_gabor1:congruency1:effector1 | 1.76 | -1.96 – 5.48 | 1.10 | -2.90 – 5.23 | | |
| accuracy_gabor1:congruency1:effector_order1 | -0.20 | -3.87 – 3.48 | 3.32 | -0.74 – 7.38 | | |
| accuracy_gabor1:effector1:effector_order1 | 2.44 | -1.36 – 6.18 | -1.97 | -6.32 – 2.46 | | |
| congruency1:effector1:effector_order1 | -0.71 | -4.42 – 2.94 | 0.84 | -3.28 – 4.99 | | |
| accuracy_gabor1:congruency1:effector1:effector_order1 | -0.53 | -4.30 – 3.21 | 0.74 | -3.32 – 4.90 | | |
| beta | 54.00 | 43.25 – 62.86 | 136.77 | 130.59 – 143.21 | 136.57 | 128.34 – 144.61 |
| **Random Effects** | | | | | | |
| $\sigma^2$ | 3570.79 | | 6254.35 | | 8553.90 | |
| $\tau_{00}$ | 15100.64 | | 25560.21 | | 31252.28 | |
| ICC | 0.19 | | 0.20 | | 0.21 | |
| N | 24 $_{subject\_id}$ | | 24 $_{subject\_id}$ | | 24 $_{subject\_id}$ | |
| Observations | 4224 | | 4510 | | 4570 | |
| Marginal $R^2$ / Conditional $R^2$ | 0.083 / 0.279 | | 0.034 / 0.234 | | 0.012 / 0.226 | |

**Table S5.** Bayesian analyses. Predicting reaction time from accuracy, congruency, effectors, and effector order in Experiment 1 (left), Experiment 2 (middle), and Experiment 3 (right).

**Replication of the analyses with the inclusion of outliers**

In summary, when replicating statistical analyses of confidence with the inclusion of outliers (10 subjects in Experiment 1, 8 subjects in Experiment 2, no exclusion in Experiment 3), results were similar (Table S6). It suggests that even if the size of the effect of action priming on confidence is slight, it is robust. Interactions that are not shown here were not significant.

Standard errors are given in parentheses. Predictors were coded as follows: Accuracy: error = -1, correct = 1; Congruency: contralateral = -1, ipsilateral = 1; Effector: different effector = -1, same effector = 1; Block order: same effector first = -1, different effector first = 1. *P < 0.05, **P < 0.01, ***P < 0.001. Response time is mean-centered. In relationship to the motor tasks, different effector corresponds to the feet and to the middle fingers for Experiment 1 and for Experiment 2, respectively.

| | **conf** | | **conf** | |
|---|---|---|---|---|
| *Predictors* | *Odds Ratios* | *CI (95%)* | *Odds Ratios* | *CI (95%)* |
| Intercept[1] | 0.65 | 0.50 – 0.85 | 0.43 | 0.31 – 0.58 |
| Intercept[2] | 1.66 | 1.27 – 2.15 | 1.29 | 0.94 – 1.75 |
| Intercept[3] | 4.19 | 3.20 – 5.45 | 3.42 | 2.50 – 4.65 |
| accuracy gabor: accuracy gabor 1 | 1.88 | 1.54 – 2.28 | 1.39 | 1.30 – 1.49 |
| congruency1 | 0.94 | 0.91 – 0.97 | 0.95 | 0.91 – 0.99 |
| effector1 | 1.00 | 0.90 – 1.10 | 1.00 | 0.90 – 1.10 |
| effector_order1 | 0.91 | 0.70 – 1.19 | 0.86 | 0.64 – 1.17 |
| rt gabor centered | 0.64 | 0.38 – 1.10 | 0.12 | 0.07 – 0.21 |
| accuracy_gabor1:congruency1 | 1.03 | 1.00 – 1.07 | 0.99 | 0.96 – 1.03 |
| accuracy_gabor1:effector1 | 1.04 | 1.01 – 1.08 | 1.00 | 0.97 – 1.03 |
| congruency1:effector1 | 0.99 | 0.96 – 1.02 | 0.99 | 0.95 – 1.02 |
| accuracy_gabor1:effector_order1 | 0.99 | 0.82 – 1.20 | 1.04 | 0.97 – 1.11 |
| congruency1:effector_order1 | 1.03 | 1.00 – 1.07 | 1.03 | 0.99 – 1.07 |
| effector1:effector_order1 | 0.92 | 0.83 – 1.02 | 0.91 | 0.82 – 1.00 |
| accuracy_gabor1:congruency1:effector1 | 1.02 | 0.99 – 1.05 | 1.03 | 0.99 – 1.06 |
| accuracy_gabor1:congruency1:effector_order1 | 1.00 | 0.97 – 1.03 | 0.97 | 0.94 – 1.01 |
| accuracy_gabor1:effector1:effector_order1 | 1.04 | 1.01 – 1.08 | 0.98 | 0.95 – 1.02 |
| congruency1:effector1:effector_order1 | 1.01 | 0.98 – 1.04 | 1.00 | 0.97 – 1.03 |
| accuracy_gabor1:congruency1:effector1:effector_order1 | 0.99 | 0.96 – 1.02 | 1.00 | 0.97 – 1.03 |
| **Random Effects** | | | | |
| $\sigma^2$ | 0.17 | | 0.20 | |
| $\tau_{00}$ | 1.08 | | 0.98 | |
| ICC | 0.13 | | 0.17 | |
| N | 26 $_{subject\_id}$ | | 32 $_{subject\_id}$ | |
| Observations | 6864 | | 5992 | |
| Marginal $R^2$ / Conditional $R^2$ | 0.206 / 0.434 | | 0.173 / 0.450 | |

**Table S6.** Experiment 1 (first conf column) & 2 (second conf column). Replication of statistical analyses of confidence as a function of accuracy, congruency, effector, effector order and their interaction with the inclusion of outliers (10 subjects in Experiment 1 and 8 in Experiment 2). Number of subjects: 26 (Experiment 1) and 32 (Experiment 2). Number of observations: 6864 (Experiment 1) and 5992 (Experiment 2).

| | acc gabor num | | acc gabor num | |
|---|---|---|---|---|
| *Predictors* | *Odds Ratios* | *CI (95%)* | *Odds Ratios* | *CI (95%)* |
| Intercept | 2.07 | 1.55 – 2.79 | 2.58 | 2.10 – 3.18 |
| congruency1 | 0.99 | 0.93 – 1.04 | 0.94 | 0.89 – 1.00 |
| effector1 | 1.07 | 0.93 – 1.22 | 1.00 | 0.85 – 1.17 |
| effector_order1 | 0.91 | 0.69 – 1.21 | 1.06 | 0.86 – 1.31 |
| rt gabor centered | 1.52 | 0.88 – 2.63 | 0.25 | 0.15 – 0.42 |
| congruency1:effector1 | 1.02 | 0.96 – 1.08 | 1.00 | 0.94 – 1.06 |
| congruency1:effector_order1 | 1.03 | 0.98 – 1.09 | 1.00 | 0.94 – 1.07 |
| effector1:effector_order1 | 0.94 | 0.83 – 1.08 | 1.04 | 0.89 – 1.23 |
| congruency1:rt_gabor_centered | 0.95 | 0.67 – 1.36 | 1.02 | 0.75 – 1.37 |
| effector1:rt_gabor_centered | 1.03 | 0.69 – 1.53 | 1.27 | 0.91 – 1.78 |
| effector_order1:rt_gabor_centered | 0.56 | 0.33 – 0.99 | 0.80 | 0.48 – 1.36 |
| congruency1:effector1:effector_order1 | 1.00 | 0.95 – 1.06 | 0.98 | 0.92 – 1.04 |
| congruency1:effector1:rt_gabor_centered | 1.21 | 0.84 – 1.72 | 1.23 | 0.92 – 1.66 |
| congruency1:effector_order1:rt_gabor_centered | 1.20 | 0.84 – 1.72 | 1.00 | 0.75 – 1.35 |
| effector1:effector_order1:rt_gabor_centered | 1.02 | 0.68 – 1.51 | 0.94 | 0.67 – 1.30 |
| congruency1:effector1:effector_order1:rt_gabor_centered | 0.80 | 0.56 – 1.14 | 0.95 | 0.70 – 1.28 |
| **Random Effects** | | | | |
| $\sigma^2$ | 0.01 | | 0.01 | |
| $\tau_{00}$ | 0.22 | | 0.20 | |
| ICC | 0.02 | | 0.03 | |
| N | 26 $_{\text{subject\_id}}$ | | 32 $_{\text{subject\_id}}$ | |
| Observations | 6864 | | 5992 | |
| Marginal $R^2$ / Conditional $R^2$ | 0.012 / 0.088 | | 0.024 / 0.080 | |

**Table S7.** Experiment 1 (first conf column) & 2 (second conf column). Replication of statistical analyses of accuracy as a function of congruency, effector, effector order and their interaction with the inclusion of outliers (10 subjects in Experiment 1 and 8 in Experiment 2). Number of subjects: 26 (Experiment 1) and 32 (Experiment 2). Number of observations: 6864 (Experiment 1) and 5992 (Experiment 2).

| | rt gabor*1000 | | rt gabor*1000 | |
|---|---|---|---|---|
| *Predictors* | *Estimates* | *CI (95%)* | *Estimates* | *CI (95%)* |
| Intercept | 438.40 | 403.55 – 472.72 | 719.54 | 691.14 – 749.15 |
| accuracy gabor: accuracy gabor 1 | 5.46 | 0.78 – 10.36 | -17.92 | -24.62 – -11.65 |
| congruency1 | 0.14 | -4.13 – 4.33 | -11.44 | -16.45 – -6.43 |
| effector1 | -2.56 | -11.31 – 6.44 | -14.06 | -29.28 – 1.40 |
| effector_order1 | 1.38 | -32.82 – 37.14 | 30.09 | 0.98 – 58.82 |
| accuracy_gabor1:congruency1 | -0.06 | -3.14 – 3.12 | -0.76 | -4.56 – 2.97 |
| accuracy_gabor1:effector1 | -0.16 | -3.35 – 3.06 | 1.54 | -2.44 – 5.55 |
| congruency1:effector1 | -1.76 | -4.92 – 1.29 | -2.57 | -6.22 – 1.01 |
| accuracy_gabor1:effector_order1 | -5.00 | -9.73 – -0.23 | -7.53 | -13.85 – -1.09 |
| congruency1:effector_order1 | -0.85 | -5.24 – 3.42 | -3.14 | -8.20 – 1.72 |
| effector1:effector_order1 | -35.55 | -45.02 – -26.32 | -9.48 | -24.10 – 6.49 |
| accuracy_gabor1:congruency1:effector1 | 2.58 | -0.53 – 5.66 | 0.62 | -3.21 – 4.24 |
| accuracy_gabor1:congruency1:effector_order1 | 0.95 | -2.11 – 4.00 | 0.79 | -2.96 – 4.49 |
| accuracy_gabor1:effector1:effector_order1 | 0.98 | -2.20 – 4.19 | -1.62 | -5.55 – 2.21 |
| congruency1:effector1:effector_order1 | -0.52 | -3.62 – 2.56 | 1.46 | -2.29 – 5.21 |
| accuracy_gabor1:congruency1:effector1:effector_order1 | -2.13 | -5.25 – 1.01 | -0.75 | -4.50 – 2.97 |
| beta | 76.17 | 70.04 – 82.05 | 141.27 | 135.34 – 147.21 |
| **Random Effects** | | | | |
| $\sigma^2$ | 6303.71 | | 6952.27 | |
| $\tau_{00}$ | 17379.07 | | 28894.25 | |
| ICC | 0.27 | | 0.19 | |
| N | 26 $_{subject\_id}$ | | 32 $_{subject\_id}$ | |
| Observations | 6864 | | 5992 | |
| Marginal $R^2$ / Conditional $R^2$ | 0.070 / 0.337 | | 0.043 / 0.240 | |

**Table S8.** Experiment 1 (first rt column) & 2 (second rt column). Replication of statistical analyses of response times as a function of accuracy, congruency, effector, effector order and their interaction with the inclusion of outliers (10 subjects in Experiment 1 and 8 in Experiment 2). Number of subjects: 26 (Experiment 1) and 32 (Experiment 2). Number of observations: 6864 (Experiment 1) and 5992 (Experiment 2).

### EEG analyses and results

**P300**. To analyze the P300 component (Figure S8), Gabor-locked epochs were corrected with a baseline of 200 ms prior to Gabor onset (Polich, 2007) at electrode Pz from 350-500 ms post-stimulus (Rausch et al., 2020). Since the P3 extended over a longer period in our study, we also investigated a larger window, 200-800 ms, which did not change the interpretation of the data.

**ERN**. We also analyzed the Error Related Negativity (ERN) component, which is observed in frontal and central electrode sites from about 50 to 300 ms after that the participant detects an incorrect choice (Scheffers & Coles, 2000). However, we did not observe an ERN component in our study (Figure S8). This was not surprising since an ERN is mainly observed in error-related paradigms (Kamarajan, 2019) and that recent studies also showed that only the Pe and not the ERN are predictive of graded changes in confidence across trials (Boldt & Yeung 2015).

**Oscillation Analyses**. We performed time-frequency analyses using Morlet wavelets. We focused on the frequencies from 4 to 7 Hz in a time period going from -1900 ms to 2500 ms time-locked at Gabor onset. We expected to observe a difference in the power of theta oscillations specifically within the first 500 ms after the onset of the Gabor since changes in this time window have been associated with cognitive control (Senoussi et al. 2022). We initially expected observing higher frontal theta activity for incongruent compared to congruent responses, since these oscillations have been associated with cognitive control (Desender et al., 2018). However, no modulation of congruency on this frequency band was found. This null result could be attributed to our task and to the nature of the conflict present in incongruent trials. Previous research showing a relationship between theta oscillations and cognitive control used conflict paradigms such as Stroop, Flankers, or Simon tasks (Bugg et al., 2008; Cavanagh & Frank, 2014), where the stimuli themselves elicited strong motor and cognitive conflict. In contrast, our task may have involved a type of conflict that might not be tracked by theta oscillations. In fact, unlike tasks such as the Stroop, where the stimulus itself prompts two or more conflicting responses, the conflict in our task was primarily between the pre-planned motor action and the required response to the Gabor. Therefore, it is plausible that the level of action inhibition and control required in our task may not have been as pronounced as in traditional conflict paradigms. The same reasoning can be applied to the null finding regarding the post-stimulus N2 component. In fact, the N2 has also been associated with response inhibition when there is an obvious conflict such as Flankers tasks or go/go-no tasks (Heil et al. 2000). Finally, we also analyzed central beta oscillations as an additional measure of motor preparation. However, it did not yield any additional information than LRP and so there are not reported in the manuscript.

**Post-response ERPs.** We performed additional analyses on post-response ERPs. We performed a cluster-based permutation test comparing low confidence (1 & 2) and high confidence (3 & 4) levels in the temporal window going from 0 ms to 600 ms after the perceptual response at Pz (see Figure S6, bottom left panel). The results yielded two significant temporal clusters: one from 288 ms to 362 ms (p = 0.04) and one from 370 ms to 600 ms (p = 0.002). Then, we performed a repeated measure ANOVA on these two clusters based on confidence & congruency, while averaging the signals from electrodes Pz, POz, CPz, P1, and P2. We averaged the activity of these electrodes rather than analyzing only brain activity at Pz channel, as this may reduce the potential impact of noise, and could highlight a possible interaction between congruency and confidence. These analyses showed very similar results to those reported in the manuscript. We observed a main effect of congruency on the temporal cluster going from 288 to 362 ms after the response, and a significant main effect of confidence on the time cluster going from 288 to 362 ms, and from 370 to 600 ms after the response. No interaction between confidence and congruency was observed.

Mean activity of Pz, POz, CPz, P1, and P2

Cluster 1 from 0.288 s to 0.362 s based on confidence

| | F Value | Num DF | Den DF | Pr > F |
|---|---|---|---|---|
| **confidence** | **7.1787** | **1.0000** | **23.0000** | **0.0134** |
| **congruency** | **5.2644** | **1.0000** | **23.0000** | **0.0312** |
| confidence:congruency | 0.0090 | 1.0000 | 23.0000 | 0.9254 |

Cluster 2 from 0.370 s to 0.600 s based on confidence

| | F Value | Num DF | Den DF | Pr > F |
|---|---|---|---|---|
| **confidence** | **16.8575** | **1.0000** | **23.0000** | **0.0004** |
| congruency | 2.9899 | 1.0000 | 23.0000 | 0.0972 |
| confidence:congruency | 0.4142 | 1.0000 | 23.0000 | 0.5262 |

We also analyzed performed a cluster-based permutation test comparing congruent and incongruent trials on the average activity of electrodes Pz, POz, CPz, P1, and P2 to check if results were different than for electrode Pz alone. We then performed a repeated measure ANOVA of the two significant time cluster observed and reported here below. The results yielded identical results as those reported in the manuscript (see Figure S6).

Mean activity of Pz, POz, CPz, P1, and P2

Cluster 1 from 0.07 s to 0.24 s based on congruency

| | F Value | Num DF | Den DF | Pr > F |
|---|---|---|---|---|
| confidence | 0.0231 | 1.0000 | 23.0000 | 0.8805 |
| **congruency** | **10.6845** | **1.0000** | **23.0000** | **0.0034** |
| confidence:congruency | 0.2233 | 1.0000 | 23.0000 | 0.6410 |

Cluster 2 from 0.26 to 0.32 s based on congruency

| | F Value | Num DF | Den DF | Pr > F |
|---|---|---|---|---|
| **confidence** | **5.0534** | **1.0000** | **23.0000** | **0.0345** |
| **congruency** | **8.7109** | **1.0000** | **23.0000** | **0.0072** |
| confidence:congruency | 0.3682 | 1.0000 | 23.0000 | 0.5499 |

=====================================================

Finally, we performed repeated measure ANOVA on each of the 5 electrodes independently. The results were very similar to those reported above and in the manuscript.

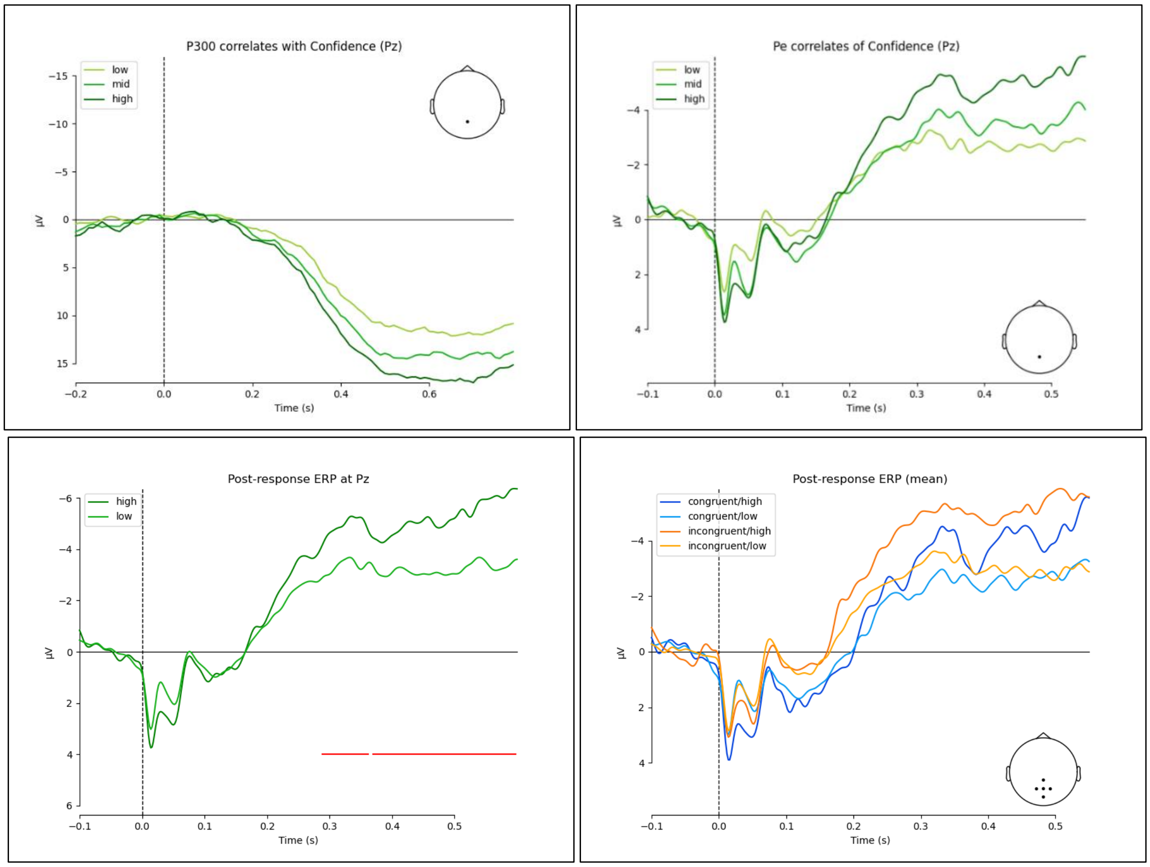


**Figure S6.** The graph in the top left panel depicts post-stimulus ERPs at Pz, time-locked at the onset of the Gabor (vertical dotted line) as a function of low (1), intermediate (2), and high (3 & 4) confidence levels. It can be observed a gradual modulation of the P300 as function of confidence. The graph in the top right panel depicts post-response ERPs at Pz, time-locked to response onset (vertical dotted line), for low (1), intermediate (2), and high (3 & 4) confidence levels. The graph in the bottom left panel depicts post-response ERPs at Pz as a function of low (1 & 2) and high (3 & 4) confidence levels (see analyses above). The horizontal red lines correspond to significant temporal clusters based on cluster-based permutation tests. The graph in the bottom right panel depicts post-response ERPs at Pz, CPz, POz, P1, P2 as a function of congruency and confidence (low -1 & 2- and high -3 & 4- confidence levels).

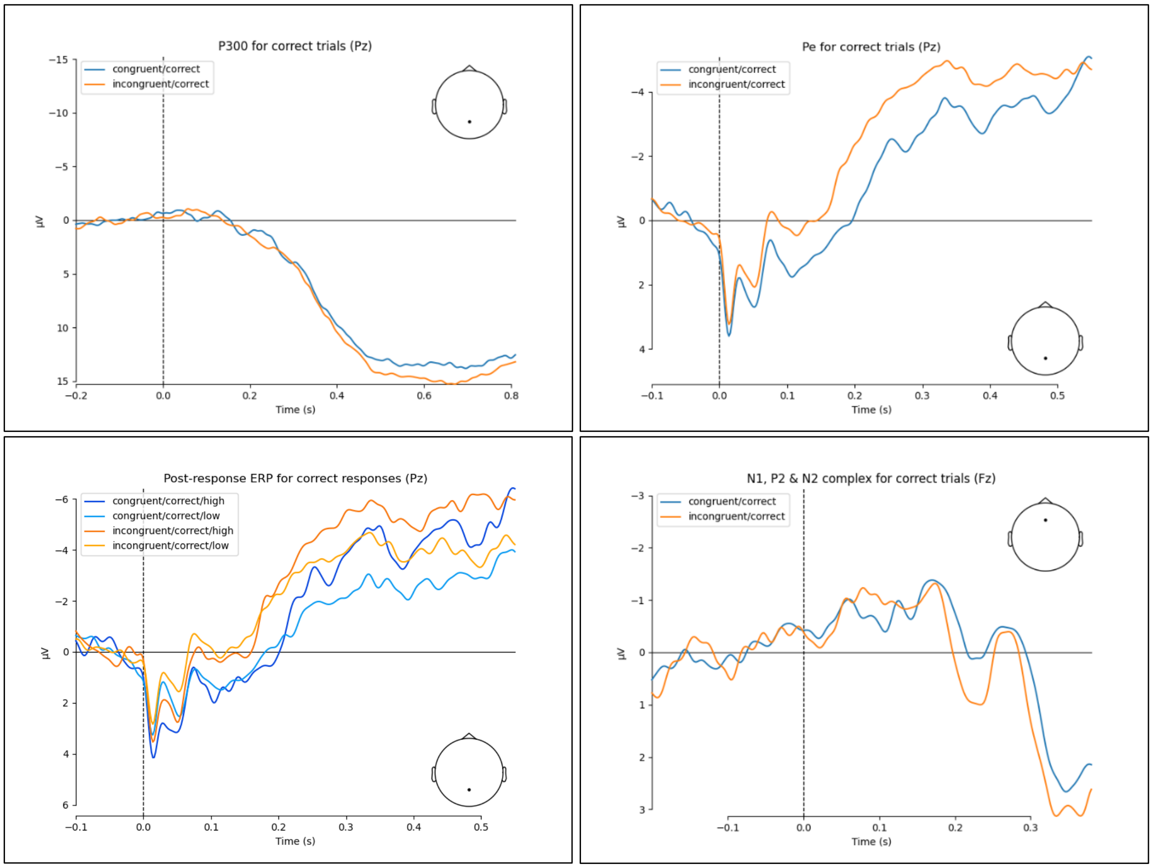


**Figure S7.** The graphs depict post-response and post-stimulus ERPs as a function of congruency (and confidence, bottom left panel) for correct responses only. In order it can be seen: post-stimulus P300 (top left panel); post-response Pe (top right panel and bottom left panel); and post-stimulus P2 (bottom right panel).

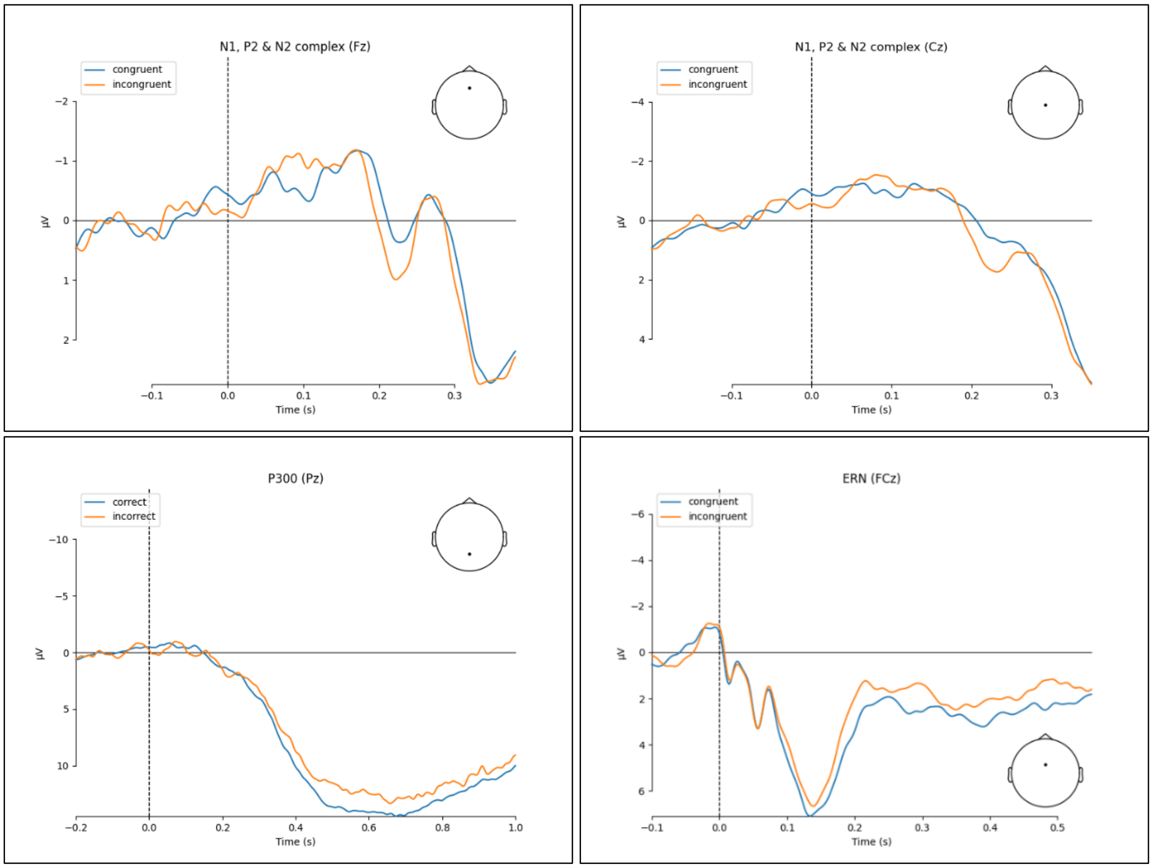


**Figure S8.** The graph in the top left panel depicts post-stimulus P2 at Fz as a function of congruency. The graph in the top right panel depicts post-stimulus P2 at Cz as a function of congruency. The graph in the middle left panel depicts post-stimulus P300 for correct and incorrect responses. The graph in the middle right panel depicts post-response ERP at FCz for congruent and incongruent responses. No Error Related Negativity (approximately 100 ms after the response) was observed in this experiment.
